# Towards a General Approach for Bat Echolocation Detection and Classification

**DOI:** 10.1101/2022.12.14.520490

**Authors:** Oisin Mac Aodha, Santiago Martínez Balvanera, Elise Damstra, Martyn Cooke, Philip Eichinski, Ella Browning, Lia Gilmour, Agata Staniewicz, Michel Barataud, Katherine Boughey, Roger Coles, Giada Giacomini, M. Cristina Mac Swiney G., Martin K. Obrist, Stuart Parsons, Thomas Sattler, Kate E. Jones

## Abstract

1. Acoustic monitoring is a scalable approach for assessing bat populations, yet automating the detection and classification of bat echolocation calls remains challenging, particularly in data-scarce regions. Although deep learning (DL) is increasingly applied to this task, most existing approaches repurpose computer-vision architectures and generate a single prediction per spectrogram clip, offering limited robustness to variable background noise and potentially constraining generality across regions and species assemblages.
2. Here, we develop BatDetect2, an open-source DL pipeline for the joint detection and classification of bat echolocation calls. BatDetect2 builds on a 2D convolutional architecture and incorporates two targeted modifications: (i) a temporal self-attention layer designed to capture long-range structure across call sequences, and (ii) convolutional layers augmented with frequency coordinates to explicitly encode frequency information directly. We evaluate model generality using five diverse datasets from four different regions: UK, Mexico, Australia, and Brazil, and conduct ablation analyses using a UK dataset spanning 17 bat species. We further assess whether a trained model can detect echolocation calls from species absent from the training data.
3. BatDetect2 consistently outperforms a traditional call-parameter extraction baseline across all datasets and evaluation metrics. Ablation analyses show that the inclusion of temporal self-attention yields a substantial species classification performance gain, increasing mean Average Precision (mAP) from 0.83 to 0.88, while frequency-coordinate augmentation provides no measurable benefit. When applied to novel species assemblages without retraining, model detection performance varies across datasets, with Average Precision ranging from 0.60 to 0.98.
4. Overall, BatDetect2 demonstrates strong and transferable performance across acoustically and taxonomically diverse regions. By jointly detecting and classifying all bat calls present in each input clip, the pipeline provides a practical and extensible tool for passive acoustic monitoring. The full training pipeline and a pretrained UK model are released through the open-source Python package `batdetect2`, enabling practitioners to develop and deploy models using their own data.

## 1 INTRODUCTION

Bats are vital bioindicators for assessing the impacts of climate change and habitat loss (Jones et al., 2009), yet significant knowledge gaps exist regarding the status of their populations (Frick et al., 2019). Acoustic monitoring, leveraging the use of echolocation by bats for navigation and prey detection (Jones and Siemers, 2011; Prat et al., 2016), offers a scalable, non-invasive and cost-effective solution for studying their activity (Gibb et al., 2018). Considerable research effort has been dedicated to automating the detection (locating calls within recordings) and classification (species identification) of bat echolocation calls in audio recordings (Zamora-Gutierrez et al., 2021), with methods evolving from the use hand-crafted acoustic features (Obrist and Boesch, 2018; Parsons and Jones, 2000; Redgwell et al., 2009) to more recent Deep Learning (DL) approaches (Mac Aodha et al., 2018; Vogelbacher et al., 2023; Khalighifar et al., 2022). However, the practical scope of existing tools remains limited, restricted to specific species or regions, and frequently inhibited by proprietary restrictions that impede transparency and accessibility. Furthermore, development is often constrained by the need for extensive datasets and specialised technical expertise (Stowell, 2022). Therefore, creating and understanding efficient methodologies for developing accurate bat detection and classification tools adequate for smaller datasets is crucial to facilitate broader bat monitoring and research (Russo et al., 2021).

Achieving accurate detection and classification of bat echolocation calls is challenging because bat calls are complex and varied. This variability stems from species-specific, regional, and habitat-dependent call characteristics (Walters et al., 2013; Montauban et al., 2021; Russo et al., 2018), which is further complicated by background noise and overlapping vocalisations from other species (e.g., small mammals and insects) (Stowell, 2022). The use of hand-crafted acoustic features, commonly referred to as call parameters in the bat literature, with traditional machine learning methods like Discriminant Function Analysis (Parsons and Jones, 2000), Random Forest (Zamora-Gutierrez et al., 2021; Bas et al., 2017; Roemer et al., 2021) or shallow Neural Networks (Walters et al., 2012; Redgwell et al., 2009) often struggle to adapt to this variability and to discriminate between similar-sounding species (Russo et al., 2018). In contrast, DL models can leverage more detailed inputs like spectrograms, potentially capturing overlooked but informative acoustic features. However, the numerous parameters that allow DL models to learn complex patterns also heighten the risk of overfitting, particularly with limited training data (Pichler and Hartig, 2023). Despite this potential for overfitting, DL models have achieved considerable success even with modest training datasets in broader ecological monitoring (Christin et al., 2019) and bioacoustics (Stowell, 2022). Nevertheless, the limited availability of data for most species and regions calls for careful consideration of how to balance model complexity with generalisation performance, and which DL architectures are best suited to address the unique characteristics of bat echolocation calls.

To date, all DL architectures applied to bat call detection and classification are adapted from the field of computer vision (Mac Aodha et al., 2018; Chen et al., 2020; Kobayashi et al., 2021; Zualkernan et al., 2020; Paumen et al., 2021; Dierckx et al., 2022; Schwab et al., 2022; Tabak et al., 2022; Yılmaz et al., 2022; Alipek et al., 2023; Brinkløv et al., 2023; Fundel et al., 2023; Vogelbacher et al., 2023; Zinck, 2023; Mahbub et al., 2024; Martinez-de-Pison et al., 2024; Meramo et al., 2025; Silva and Herrera, 2026). Specifically, 2D Convolutional Neural Networks (CNNs) have been widely employed to analyse spectrograms derived from audio recordings. In images, information needed for object identification is spatially concentrated (locality), and an object’s identity remains consistent regardless of its position (translation invariance). The ability of CNNs to exploit these inherent properties through the convolutional operation allows for a more efficient architecture with significantly fewer parameters than fully-connected networks, leading to improved performance in image classification tasks (LeCun et al., 2015; Menghani, 2023). However, these core assumptions in computer vision do not necessarily translate well to the analysis of audio data.

Bat echolocation calls exhibit unique spectro-temporal characteristics that challenge the direct application of standard CNN architectures. For instance, the distinct frequency ranges of calls emitted by different bat species, which reflect adaptations to their foraging environments and prey types (Denzinger and Schnitzler, 2013; Walters et al., 2013), imply that translation invariance may not hold in the frequency dimension for bat calls. Furthermore, as bats typically emit sequences of echolocation calls, where the inter-pulse interval and the overall temporal structure are often crucial for accurate species identification, the assumption of locality might not be entirely appropriate in the temporal dimension. While analysing short audio clips, typically shorter than 50 milliseconds, reduces the impact of this issue (Mac Aodha et al., 2018; Chen et al., 2020; Kobayashi et al., 2021; Khalighifar et al., 2022), it leaves out the potentially discriminative information encoded in the longer temporal structure of call sequences. Using CNNs to analyse longer audio clips, as done by Paumen et al. (2021), Zualkernan et al. (2020), and Tabak et al. (2022), typically requires increasing model depth and size to capture longer temporal relationships (Simonyan and Zisserman, 2015; He et al., 2016), thereby demanding substantially more training data for robust results. Additionally, using longer audio clips includes more background noise, which may confuse the learning process without specific guidance on call location and lead to degraded classification performance (Gadot et al., 2024). This naturally raises the question of how to design a model that can effectively capture both the spectral and temporal structure of bat echolocation calls while remaining compact and efficient.

Despite the recent progress in DL-based solutions for bat detection, a gap persists between the latest research advancements and the open-source tools available to practitioners. Of the DL models developed for automated bat detection, only Mac Aodha et al. (2018), Alipek et al. (2023), and Fundel et al. (2023) offer open-source implementations. Furthermore, training and using custom DL models presents several additional challenges. First, even with the availability of open-source tools for developing bioacoustic models (Lapp et al., 2023), training models typically requires proficiency in programming and machine learning, presenting a barrier for many practitioners. Secondly, training DL models to identify particular species requires substantial amounts of labelled data, which are often scarce for many bat species and regions. However, being able to detect bat calls, even without identifying their species, would already provide significant value. Finally, the “black box” nature of many DL models makes it difficult to interpret their decision-making processes. This lack of transparency is particularly problematic when a model generates a single prediction from a long audio clip containing multiple calls from different species (Dierckx et al., 2022). Given these challenges, there is a clear need for user-friendly tools that empower practitioners to develop and deploy robust DL models for bat monitoring, without requiring extensive programming or machine learning expertise.

Here, we develop BatDetect2, a novel model for bat echolocation call detection and species classification from acoustic data. BatDetect2 incorporates two key modifications to the standard CNN architecture, specifically designed to enhance the model’s ability to better capture spectral and temporal characteristics. Furthermore, the model also provides interpretable predictions that illustrate where in the input spectrogram, in terms of frequency and time, the model has detected a call. We train and evaluate BatDetect2 using five challenging datasets from four different geographical regions (UK, Mexico, Australia, Brazil), and compare its performance to existing call parameter-based methods. Using a UK dataset of 17 bat species, we evaluate the impact of these modifications on the model’s overall performance, as well as on its performance for each individual species. Finally, using the model trained on UK data, we evaluate its potential for detecting echolocation calls from other regions. To facilitate adoption and further development by practitioners, the complete pipeline, including code and trained models, is made publicly available as an open-source Python package: `batdetect2` (accessible at https://github.com/macaodha/batdetect2). This package enables users to train new models from scratch and deploy them for automated analysis of their own datasets.

## 2 MATERIALS AND METHODS

### 2.1 Acoustic event detection and classification

Within a single audio recording, multiple distinct acoustic vocalisation events (e.g., a bat echolocation call) may occur, each occupying a distinct Region of Interest (RoI) in the time–frequency domain. Our goal is to develop a model that takes an ultrasonic audio recording as input, represented as a spectrogram, and outputs a set of predictions related to the events of interest. In this work, the events of interest are bat echolocation calls, and each prediction corresponds to one call, providing its location within the spectrogram, its temporal and spectral properties, and its predicted species.

More formally, each predicted event is represented as **o** = [**g**_roi_, *p*_detection_, **p**_species_], where **g**_roi_ represents the geometry describing the RoI, *p*_detection_ is the confidence that the event is relevant (i.e., an echolocation call), and **p**_species_ is a C-dimensional vector containing the confidence score assigned to each of the C species of interest. We define the RoI using a bounding box, **g**_roi_ = [*t* _start_, *f*_low_, *t*_end_, *f*_high_], which specifies the call’s start and end time as well as its lowest and highest frequencies. Note that this representation is distinct from conventional acoustic classification models that only attempt to determine the species present in a short duration input spectrogram (Stowell, 2022). We also depart from traditional Sound Event Detection (Mesaros et al., 2021) by incorporating explicit frequency bounds (*f*_low_,*f*_high_), a characteristic particularly suitable for bat calls due to their relatively clear frequency ranges.

### 2.2 Model architecture

We implement our joint classification and detection model as a deep neural network, which we refer to as BatDetect2. Inspired by computationally efficient one-stage object detection methods from computer vision, e.g., Zhou et al. (2019), this model directly predicts the location and size of each event (i.e., echolocation call) in the input. BatDetect2 makes use of a U-Net-style architecture (Ronneberger et al., 2015), with an encoder that extracts features from the input spectrogram, followed by a decoder that generates the predicted size and location of each echolocation call along with the corresponding species’ probabilities (Fig. 1). The model uses skip connections, which allow for the propagation of higher-resolution feature information (in terms of frequency and time) from the encoder to the decoder. The full architecture is described in Table S6. Crucially, while BatDetect2 is based on the U-Net architecture, which typically employs 2D-convolutional layers, it incorporates two key modifications to better capture the distinctive spectral and temporal characteristics of audio signals described below.

**FIGURE 1.**
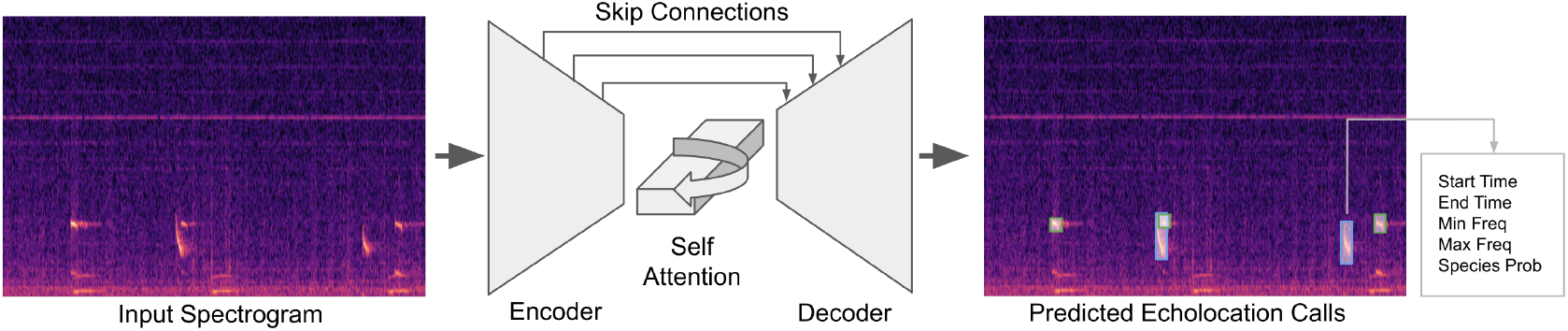
Overview of BatDetect2 architecture. The model consists of a convolutional neural network-based encoder and decoder with skip connections that share extracted features from the encoder to decoder. The encoder and decoder consists of modified convolutional layers that incorporate coordinate information. It utilises a self-attention layer in the middle of the model so that it can reason over a longer temporal scale. In contrast to most existing deep learning-based bat call classifiers, BatDetect2 directly predicts the time in file of each event of interest, along with the duration of the event, the frequency range, and the species.

To allow the model to reason about the temporal structure of the echolocation calls, we incorporate a self-attention layer into the middle of the network (Fig. 1). The self-attention mechanism, commonly associated with the Transformer architecture (Vaswani, 2017), allows the model to “attend” to information from different points in time in the audio file and has proven effective in natural language processing for capturing long-range dependencies. The network processes the fine-grained spectrogram input (1 ms per time bin) by transforming it into a coarser sequence of feature vectors (approximately 8 ms per bin). The self-attention layer operates on this coarser representation. For each time step, it calculates its relationship to all others, generating attention weights (Fig. S2). These weights are used to compute a context-aware representation that helps the model refine its estimate of a species’ presence. A decoder then uses these enhanced representations to generate final predictions at the original, fine temporal resolution. Note that, unlike vision transformers such as ViT (Dosovitskiy et al., 2021) used in Fundel et al. (2023), which employ multiple self-attention layers across both time and frequency, our model uses a single self-attention layer operating solely on the temporal dimension, resulting in significantly reduced computational burden.

To mitigate the undesirable translation invariance along the frequency axis, we use two specialised building blocks: `CoordConvDown` and `CoordConvUp`. At a high level, the `CoordConvDown` layer takes a tensor as input and returns a spatially downsized version of it as an output. Unlike standard convolutional layers, which exhibit translation invariance, `CoordConvDown` appends non-learnable, normalised coordinates along the frequency (vertical) axis of the input tensor. This modification is potentially valuable because the absolute frequency of an echolocation call provides valuable discriminative information for bat species identification. In contrast to the original `CoordConv` approach (Liu et al., 2018), coordinate information is not added along the temporal axis. This design choice preserves the desired time-translation invariance, allowing the model to recognise calls regardless of their precise position within the recording. The `CoordConvUp` layer performs the inverse operation of `CoordConvDown`, upsampling the feature maps.

For each input spectrogram, the model initially generates an intermediate “feature” map, an array with the same height and width as the input. Each pixel in the feature map encodes a 32-dimensional feature vector, representing learned acoustic characteristics at the corresponding time-frequency location. This feature map is then used to produce two primary outputs: a “class” map (*Ŷ*) and a “size” map (*Ŝ*). Both of these outputs have the same height and width as the input spectrogram.

The class map *Ŷ* indicates where the model predicts echolocation calls are located within the spectrogram and what species they belong to. Each pixel in *Ŷ* contains a vector with a length equal to the number of bat species in the dataset plus one. This additional element represents a “background” class, indicating the absence of a call. Each value in the vector represents the confidence of the model that a call of a particular species is present at that pixel’s corresponding time and frequency. Ideally, only the pixel corresponding to the bottom-left corner of a call’s bounding box should have a high confidence value for the correct species, and all other pixels should indicate “background.” The size map, *Ŝ*, provides information about the estimated size of the detected echolocation calls. Each pixel in *Ŝ* contains two values: an estimated height and width of the bounding box around any echolocation call detected at that pixel’s location.

As a final step, this output is passed through a non-maximal suppression layer, implemented via max pooling, in order to extract the local peak detections (Zhou et al., 2019). This step prevents the model from predicting multiple calls very close to each other (i.e., within a few milliseconds).

### 2.3 Audio preprocessing

To prepare the raw audio for model processing, the input audio is transformed into spectrograms as follows. Firstly, the input audio is resampled to 256 kHz using the polyphase method from `scipy` (Virtanen et al., 2020). We selected `scipy`’s polyphase resampling method for its low computational cost, making it well-suited for processing large audio datasets. We then compute the magnitude spectrogram using a Short Time Fourier Transform with a window size of 512 samples and a window overlap of 75%. As the bat echolocation calls recorded for this study are found only within a specific frequency range, we retain only the bands between 10 kHz and 120 kHz. For robustness to volume variations, the spectrogram is normalised using PCEN (Wang et al., 2017), which Lostanlen et al., 2019 showed to be a more effective frontend than traditional log-scaled spectrograms. Following Aide et al. (2013) and Mac Aodha et al. (2018), we also subtract the mean value from each frequency band to mitigate the impact of any constant background noise. Finally, we use bilinear interpolation to resize the temporal dimension down by a factor of two and resample the frequency bands into 128 bins. Consequently, a one-second input audio file results in a spectrogram of size 128 × 1024.

### 2.4 Model training

BatDetect2 is trained using a supervised learning approach, where it is provided with input spectrograms and corresponding target outputs. These target outputs are derived from manually created bounding box annotations of bat echolocation calls. The target for the “class” map *Ŷ* is constructed by creating a series of “heatmaps,” one for each species. Each species-specific heatmap is initially set to zero everywhere except in the vicinity of the bottom-left corner of each call annotated for that species. At these locations, a Gaussian kernel with a standard deviation of 2.0 is applied, creating a localised peak with smoothly decaying values, a method consistent with prior work in object detection (Zhou et al., 2019). Unlike Zhou et al. (2019), who parameterises bounding boxes using their centre point, we instead use the point corresponding to the start time and minimum frequency of each echolocation call. We chose the minimum frequency point because it exhibited less inter-species variability compared to the centre frequency across the training data, with the notable exception of the two UK *Rhinolophus* species. Once the individual species-specific heatmaps are generated, they are stacked together to form the final target map for *Ŷ*. To create the target for the “size” map *Ŝ*, the height and width of the bounding box of each annotated call are calculated in pixel units. These values are then assigned to the pixel location corresponding to the bottom-left corner of the bounding box. All other pixel locations, which correspond to areas without annotated calls, are assigned values of zero for both height and width.

The model is trained end-to-end using a three component loss function which includes a detection loss, a classification loss, and an event size loss. The detection loss is computed by comparing the predicted class map (*Y*) to the ground truth class map (*Ŷ*), considering only the complement of the background class (i.e., the sum of species-specific heatmaps). In contrast, the classification loss compares all individual species-specific heatmaps. Both losses are implemented using a focal loss, as described by Lin et al. (2017). This loss function is particularly well-suited to this scenario as most spectrogram pixels do not contain calls, resulting in a substantial class imbalance between “background” and “call” pixels. The event size loss is calculated using an L1 loss that penalises the absolute difference between predicted and actual dimensions. This loss is computed only for pixels corresponding to true calls (see SI E for a detailed description of the training losses).

During training, clips of 256 ms duration are extracted at random from the training recordings. To increase the variation in the input audio, we perform a series of data augmentations at training time. These augmentations include: random linear combination of two input audio files (Zhang et al., 2018), simulated echo, random volume scaling, temporal stretching, and time and frequency masking (Park et al., 2019). The probability that any one augmentation is applied is 0.2, and multiple augmentations can be applied simultaneously.

The model and training code are implemented in `PyTorch` (Paszke et al., 2019). We train the model end-to-end using the Adam optimiser (Kingma and Ba, 2017), starting with an initial learning rate of 0.001, a cosine annealing learning rate schedule, and a batch size of eight. Training is performed for 200 epochs with no early stopping, except for the Australia dataset, which is trained for 800 epochs.

### 2.5 Audio datasets

We train and evaluate BatDetect2 on five different full spectrum ultrasonic acoustic datasets. In preparation for training, these datasets are manually annotated to obtain bounding boxes that encompass each individual echolocation call within an audio file. To generate these annotations, an early version of `whombat` (Martínez Balvanera et al., 2024), was employed. The annotations were created primarily by the authors, with additional contributions from bat experts recognised in the acknowledgments.

Annotators were instructed to draw boxes around each individual echolocation call, irrespective of how faint the call was. Unless otherwise specified, the annotated audio files had information at the file-level related to which species were present in the recording. They then assigned the recording-level species class label to an annotation unless it differed from a prototypical echolocation call for that species. Harmonics were not annotated as part of the main call. In cases where it was not possible to assign the correct class label (i.e., species), or when multiple species were present in a file, annotators marked unknown calls as being from a generic “Bat” class.

#### 2.5.1 UK datasets

The primary dataset used in this study, which contains recordings of all 17 bat species known to breed in the UK (Fig. S3), is composed of three distinct sets of recordings: those containing known species calls, those containing unknown species calls, and empty field recordings.

The first set was collated from six distinct sources, including the Bat Conservation Trust and individual contributors, ensuring a wide variety of recording devices and acoustic environments capturing free-flying bats near known roosts. The majority of the recordings were made in the UK, but there were also some additional files included from the species of interest that were recorded elsewhere (e.g., in Europe). In total, these contain 2,809 distinct audio files with a mean duration of 1.04 seconds, encompassing 34,635 annotated echolocation calls, the majority of which have confirmed species labels. Secondly, we incorporated additional 4,225 0.384-second duration files adapted from Mac Aodha et al. (2018) and collected in the iBats Program (Jones et al., 2013). This data was recorded using Tranquility Transect detector using a time expansion factor of ten. This adds an additional 6,842 annotated bat calls that do not have a confirmed species label. Finally, we also add 345, one-second duration, urban park soundscape recordings without bat sounds from London, UK, collected using the recording devices described in Gallacher et al. (2021). These “empty” files are intended to help the model learn to better distinguish between bat calls and background noise.

We split the UK data into two different train and test sets, UK_same_ and UK_diff_. For UK_same_, we randomly assign files to the test set, ensuring a maximum of four files per species and data source. The remaining files are kept for the training set. This results in 7,010 train files and 369 test files, containing 36,955 and 4,522 calls, respectively (Table S1). UK_diff_ is a more challenging split. Here we hold-out the largest single data source for testing. This leaves 5,911 training and 1,468 test files, containing 24,315 and 17,162 echolocation calls, respectively (Table S2). This second split represents a more challenging test-case in terms of generalisation where the data is guaranteed to be very different from the training set. This also results in a reduction in the overall amount of training data, both in terms of sheer quantity but also diversity. Both the UK_same_ and UK_diff_ sets include the 4,570 files without species labels as part of their training data.

### 2.5.2 Yucatan data

The second dataset consists of 1,193 one second audio clips extracted from 285 passive acoustic recordings from the Yucatan peninsula in Mexico. The data was collected as part of a study by MacSwiney G. et al. (2008). It is smaller in size than the UK dataset, but is representative of the type of data that would be feasible to collect and annotate as part of a smaller-scale monitoring project. The annotations from the original study were used and then expanded to ensure that all audible echolocation events were annotated. The final annotated dataset contains 10,020 echolocation calls from 17 different species (Fig. S4) distributed across these 1,193 one-second clips. We divided the data into 911 training and 282 test clips, making sure to separate at the original recording-level, to ensure that clips from the same recording were not in both sets (Table S3).

#### 2.5.3 Australia data

This next dataset consists of a set of 14 bat species which can be found in the major cotton growing region on the north central plains of New South Wales and adjacent areas in central southern Queensland. Bat calls were recorded from captured individuals following positive species identification, with recordings taken sufficiently after release to capture natural acoustic behavior. This dataset features sympatric species with highly overlapping call characteristics (e.g., within the *Nyctophilus* and *Ozimops* genera), which makes acoustic classification particularly challenging (Fig. S5). The data was randomly split at the file level, with 80% of the recordings for a species staying the train set, and the rest in the test. This resulted in 4,569 and 1,327 individual calls in the train and test sets respectively (Table S4).

#### 2.5.4 Brazil data

The final dataset presents a distinct challenge as it lacks confirmed species labels. It contains 320 recordings of ten second duration, each collected between January and March 2019 in south-eastern Brazil using AudioMoth recorders (Hill et al., 2019). As the identity of recorded bat species could not be independently verified, calls could not be assigned species labels during annotation. Instead, we created three “sonotypes” based on the dominant frequency component of each call and labelled individual calls accordingly (Fig. S6). Like the other datasets, the annotation was performed manually, where the protocol again stipulated that all echolocation call instances in each recording should be annotated. We split the data into 256 train files and 64 test files, which resulted in 7,989 and 2,010 calls respectively (Table S5).

### 2.6 Model Evaluation

To evaluate model performance, we used the trained models to generate predictions on all recordings within the test sets. For each recording, the entire audio file was fed into the model at once. Recordings may vary in duration, so they are padded with silence at the end to meet the input requirements of the model (specifically, multiples of 256 ms). To quantify model performance, we use four different evaluation metrics detailed below.

Firstly, detection average precision (‘AP Det’), evaluates the ability of the model to correctly identify all echolocation calls in the test data. This metric calculates the precision and recall resulting from varying a threshold on the model output predictions for the ‘Bat’ versus ‘Not bat’ task. We then average over these different thresholds to quantify the area under the precision-recall curve, using the interpolation method used in Everingham et al. (2009). A prediction is counted as a true positive if its estimated start time is within 10 ms of the start of a ground truth echolocation call. When multiple predictions match the same ground truth call, only the prediction with the highest confidence score is designated as a true positive. All other matching predictions are counted as false positives, unless they match a different ground truth call. This is the same evaluation criteria used in Mac Aodha et al. (2018).

To assess the ability of the model to accurately assign the correct species label to each detected call, we report the mean average precision across the classes (‘mAP Class’). This involves taking the per-class average precision and then averaging this value over all classes. By design, this metric weights all classes equally, regardless of the number of calls for each class in the test set. For this calculation, the match preference is based on the class confidence score, and we exclude any matched calls for which the ground truth species is unknown (i.e., the generic ‘Bat’ class).

‘mAP Class’ suffers from one major limitation. As the classes are evaluated independently, it does not highlight cases where the underlying model may be poorly calibrated and thus require different output thresholds for each class. Calibration issues like this can result from class-level data imbalances in the training data. To overcome this limitation, we also report a third precision based metric which we refer to as ‘Top Class’. Here we simply take the top predicted class label, along with its corresponding probability, for each detected call and then evaluate the average precision as above. In this case, a prediction is counted as a true positive if it matches a ground truth call and it’s the top predicted class matches the ground truth species. Here, match preference is based on the top predicted class confidence score. As with ‘mAP Class’, matches are excluded if the ground truth species is unknown. Unlike ‘mAP Class’, this metric can be biased if there is a large imbalance in the classes in the test set.

The final metric, ‘mAP File’, evaluates the file-level classification performance. This metric assesses the ability of the model to accurately predict the presence or absence of each species within a given test file. Notice that, in contrast to a call which can only be produced by a single species, multiple species can occur in a single file, making this a multi-label task. To obtain the file-level confidence score for each class from the multiple individual call predictions within a given file, we take the maximum confidence score for that class among all detections within the file. The confidence score is set to 0 if no detections for that class are present in the file. We then compute the average precision at the file level for each class and report the mean across all classes. The best possible score for each of these four metrics is 1.0, and the worst is 0.0.

### 2.7 Experiments

#### 2.7.1 Evaluating performance across diverse settings

To evaluate the generality of the proposed pipeline, we compare its performance across diverse settings to a traditional bat call parameter extraction baseline. Both the BatDetect2 model and the baseline are trained from scratch on all five datasets (UK_same_, UK_diff_, Yucatan, Brazil, and Australia) using identical training and testing data and are evaluated using the same metrics. Both models are trained three times on each dataset to account for stochastic fluctuations in the training process, and the final results are averaged.

The baseline is a Random Forest (Breiman, 2001) classifier trained on acoustic features extracted by the Tadarida-D model (Bas et al., 2017). Tadarida-D consists of two main components: (i) a generic sound event detector and (ii) a feature extractor. We chose to use Tadarida-D since it is the only open-source pipeline that is able to do detection of bat echolocation events instead of clip-level prediction of species presence/absence. The extracted detections and their corresponding acoustic features are then used to train the Random Forest classifier to distinguish true echolocation calls from background noise and assign species labels (see SI F.1 for details).

This baseline allows for a controlled comparison with a traditional call parameter-based method by using the same audio data and ground truth annotations for both training and evaluation. However, it is important to emphasise that while we are using Tadarida-D, the baseline is *not* directly equivalent to the full Tadarida method as we do *not* make use of their pre-trained models, labeling interface, classification code, or post-processing steps.

#### 2.7.2 Evaluating the impact of architectural modifications

To assess the impact of the proposed architectural modifications on model performance, we conducted an ablation study. In addition to the full BatDetect2 model, we trained and evaluated three variants: (1) NoSelfAttn, which removes the self-attention layer; (2) NoCoord-Conv, which replaces `CoordConv` layers with standard convolutions; and (3) FullyConv, which removes both modifications. All four models are trained and evaluated on the more diverse UK_same_ dataset split using identical training and evaluation protocols. Again, each model is trained three times and the results are averaged.

To analyse performance differences at the species level, we compute per-class average precision each species in the UK_diff_ dataset, in addition to the standard global evaluation metrics. This allows for a more granular analysis of how each architectural modification affects the detection and classification accuracy for individual bat species.

#### 2.7.3 Evaluating detection in novel regions

While BatDetect2 can only classify species it was trained on, it may still be able to detect echolocation calls from species it has never encountered if they are acoustically similar to the training data. We tested this generalisation ability by applying the model pretrained on the UK_same_ dataset directly to the Yucatan, Brazil, and Australia datasets without modification or further training. We refer to this UK-trained model as BatDetect2_UK_. The UK_same_ dataset was chosen as the pre-training source for its large size and diversity.

This off-the-shelf detection performance was then compared against three baselines. Firstly, we used the Random Forest (RF_UK_) baseline as described previously, also trained on the same UK_same_ dataset and applied to the target datasets without further training. Secondly, we included the off-the-shelf Tadarida-D detector, used without modification. Finally, we compared these against a simple, generic detector (Onset) based on the acoustic strength envelope (see SI F.2 for details). As this evaluation focuses solely on detection, we report the AP Det metric and supplement it with the F1-score. For the F1-score, we computed recall and precision at an illustrative confidence threshold of 0.5, when appropriate. Since the Onset and Tadarida-D baselines do not output a confidence score, AP Det was not computed for them. Both BatDetect2_UK_ and RF_UK_ were trained three times on the UK_same_ dataset, evaluated on the target dataset, and the results were then averaged.

## 3 RESULTS

### 3.1 Detection and classification performance

BatDetect2 shows a substantial improvement over the Random Forest baseline across all datasets and evaluation metrics (Table 1). The performance difference, measured by mean average precision (mAP Class), ranges from 0.05 to 0.33 across the datasets. While the Random Forest baseline achieves reasonable performance on the comparatively less complex Brazil dataset, it exhibits significantly lower performance on the remaining datasets. The detection performance of BatDetect2, as indicated by the AP Det metric, shows consistent results above 0.91 in all cases (Table 1). BatDetect2 appears to be robust to background noise, as even in the presence of repetitive high-frequency noise, or sudden broad band clicks, the model does not produce false positives (Fig. 2).

**TABLE 1.** Performance of BatDetect2 model (BD2) compared to the Random Forest baseline (RF) with traditional bat echolocation call features. Both models are evaluated using the same five test datasets. For each of the metrics, higher numbers are better, and the results are averaged over three runs. For each metric and dataset, the best score is highlighted in bold.

| Dataset | AP Det |  | mAP Class |  | Top Class |  | mAP File |  |
| --- | --- | --- | --- | --- | --- | --- | --- | --- |
|  | BD2 | RF | BD2 | RF | BD2 | RF | BD2 | RF |
| UK<sub>same</sub> | **0.970** | 0.881 | **0.885** | 0.712 | **0.835** | 0.637 | **0.914** | 0.857 |
| UK<sub>diff</sub> | **0.964** | 0.902 | **0.813** | 0.584 | **0.691** | 0.466 | **0.816** | 0.678 |
| Yucatan | **0.919** | 0.645 | **0.811** | 0.448 | **0.803** | 0.480 | **0.778** | 0.541 |
| Australia | **0.978** | 0.952 | **0.772** | 0.623 | **0.692** | 0.531 | **0.854** | 0.752 |
| Brazil | **0.918** | 0.873 | **0.970** | 0.915 | **0.947** | 0.913 | **0.919** | 0.904 |

**FIGURE 2.**
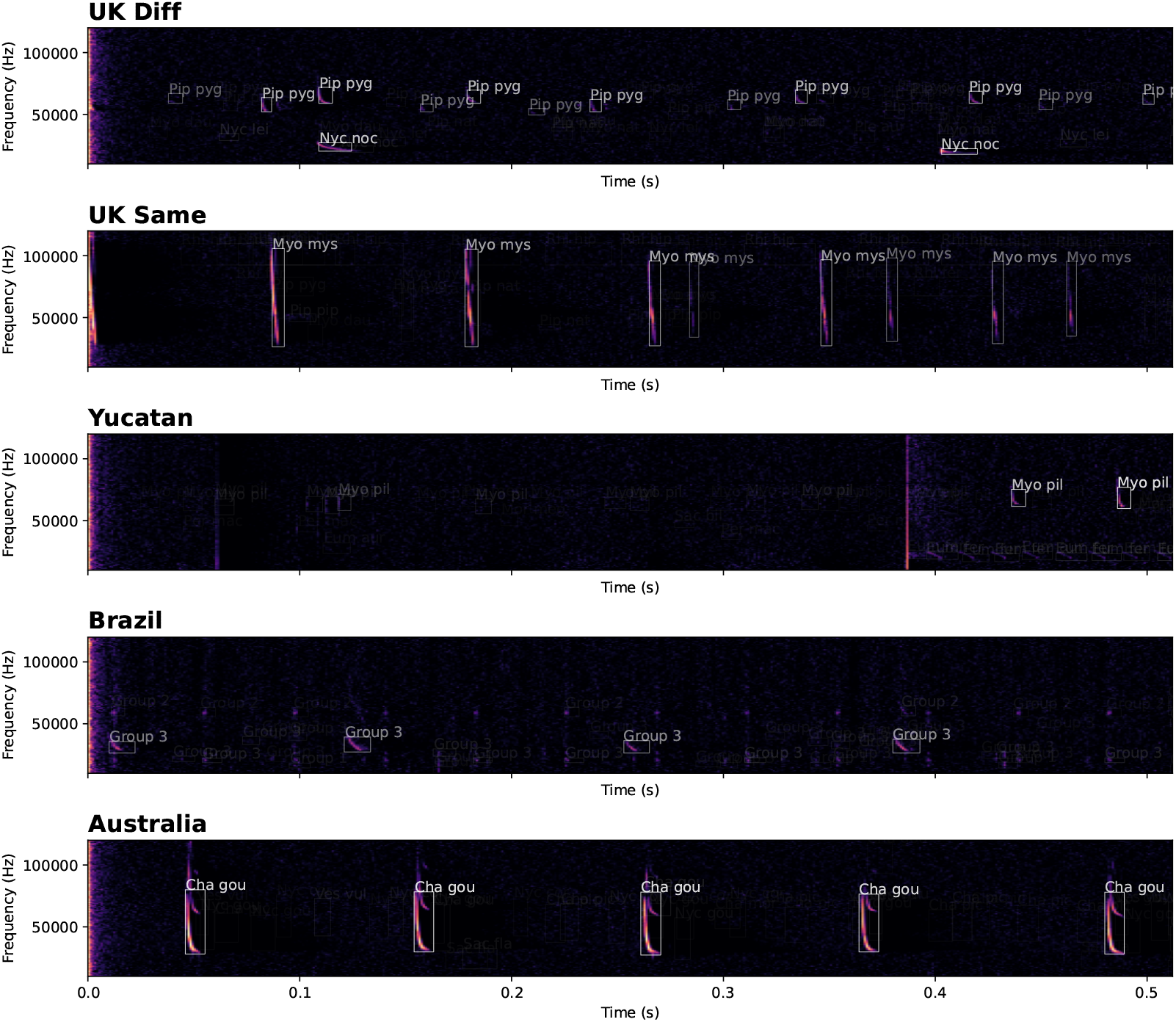
Predictions from the BatDetect2 model. Each row represents a different audio file selected from the test sets of the UK_same_, UK_diff_, Yucatan, Brazil, and Australia datasets, ordered from top to bottom. The intensity of an individual predicted bounding box indicates the model’s confidence, with a brighter white value indicating more confident. The text above each box corresponds to the highest scoring class label.

On the contrary, the call-level classification performance, measured by mean average precision (mAP Class) and top class average precision (Top Class), showed considerable variability across datasets, ranging from 0.77 mAP Class and 0.69 Top Class for the Australia dataset to 0.97 mAP Class and 0.95 Top Class for the Brazil dataset. The performance discrepancy between the UK_diff_ and UK_same_ datasets likely reflects the more challenging train-test split employed in the former, coupled with the associated reduction in the size of the training data. This split results in a test set that is less similar to the training data, despite covering the same species, thus posing a greater challenge for generalisation. Classification performance on the Yucatan dataset is similar to that on the UK_diff_ dataset, whereas the Random Forest baseline exhibits a substantial drop. In contrast, both models show a noticeable performance decrease on the Australia dataset, partially explained by challenging overlapping species and fewer distinct training files. Conversely, performance on the Brazilian dataset remains very high, largely due to fewer overall classes and the grouping of acoustically similar calls into broader sonotypes.

Examining the performance of the BatDetect2 model on the UK_same_ dataset split at the species level shows challenges in accurately classifying certain species (Fig. 3A). Notably, the average precision for *Myotis bechsteinii, Myotis brandtii*, and *Plecotus auritus* remains below 0.8. While *Nyctalus leisleri* and *Plecotus auritus* exhibit precision above 0.8 at a recall of 0.6, their performance declines at higher recall values. In contrast, all other species achieve a precision of at least 0.8 at a recall of 0.8. When comparing the top-predicted species with the true species we see considerable inter-species confusion within the *Myotis* genus (Fig. 3B), where calls from both *Myotis bechsteinii* and *Myotis brandtii* are frequently mispredicted as *Myotis daubentonii*. When excluding the poorly-performing *Myotis* species, no correlation between the number of training examples and average precision is apparent.

**FIGURE 3.**
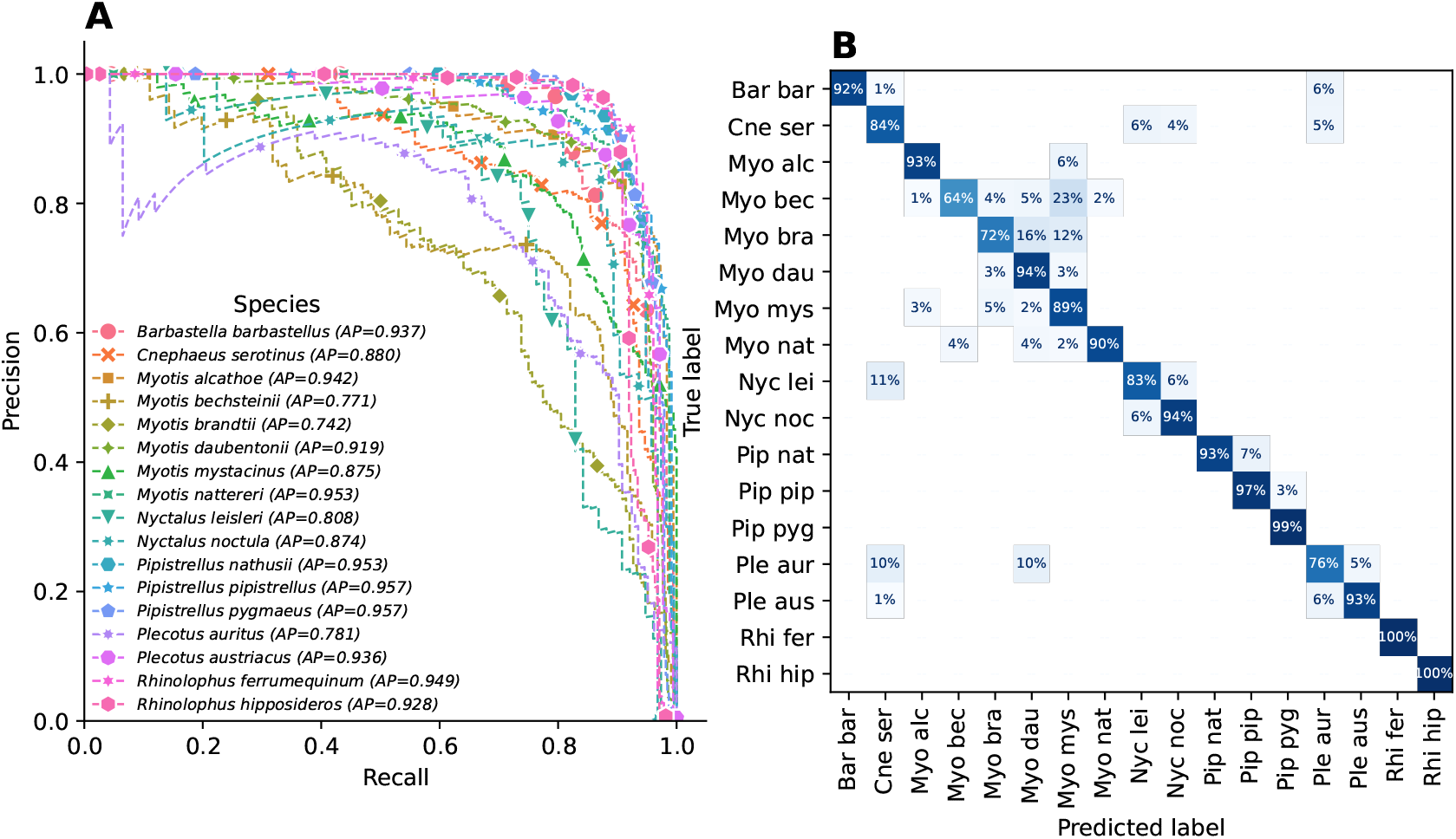
Per-species performance of BatDetect2 on the UK_same_ dataset. (A) Precision-recall curves for each species in the UK_same_ dataset. (B) Call-level confusion matrix for the UK_same_ dataset. The matrix compares the true species against the top-predicted species for all matched echolocation call events. A prediction and ground truth annotation are matched if their start times are within 10ms of each other. We exclude matches where the true species is unknown, as well as unmatched predictions or ground truth calls (i.e., false positives and false negatives). Predictions with a class score below 0.2 are also excluded. Rows are normalised to sum to 100% and values below 1% are omitted for clarity.

It takes BatDetect2 just under four minutes to process and save the results for 424, ten second duration, 384 kHz AudioMoth recordings using a GPU, i.e., 70.6 minutes of ultrasonic audio data in total. Tadarida-D, which does not utilise a GPU, takes 2.5 minutes for detection and feature extraction for the same data. Note that this processing time does not include the evaluation of the Random Forest. This benchmarking was performed on a workstation which contained an Intel i7-6850K CPU and an Nvidia TITAN Xp GPU.

### 3.2 Impact of architectural modifications

We found that including the self-attention mechanism in the model resulted in a substantial improvement in performance (Table 2). Specifically, while call detection (measured by AP Det) remained unaffected, classification accuracy improved considerably. The mAP Class and Top Class metrics increased markedly from 0.84 to 0.89 and from 0.78 to 0.84, respectively, upon inclusion of the self-attention layer. On the contrary, the inclusion of coordinate convolutional layers only yielded a small yet positive impact on performance.

**TABLE 2.** Performance of BatDetect2 variants on the UK_same_ test set. Metrics are the average precision for detection (AP Det), the mean average precision for classification (mAP Class), the top class average precision (Top Class), and the file mean average precision (mAP File). The ‘NoSelfAttn’ variant is identical to the full BatDetect2 model but omits the self-attention layer. The ‘NoCoordConv’ variant is identical to the full model but does not incorporate frequency coordinate information into the convolutional layers. Finally, the ‘FullyConv’ variant is a baseline model that does not incorporate any of the above modifications. Each model was trained three times on the UK_same_ training set, and the mean performance on the corresponding test set is reported.

| Model | Self-Attention | CoordConv | AP Det | mAP Class | Top Class | mAP File |
| --- | --- | --- | --- | --- | --- | --- |
| BatDetect2 | ✓ | ✓ | 0.970 | 0.885 | 0.835 | 0.914 |
| NoSelfAttn |  | ✓ | 0.966 | 0.835 | 0.780 | 0.893 |
| NoCoordConv | ✓ |  | 0.962 | 0.878 | 0.824 | 0.917 |
| FullyConv |  |  | 0.967 | 0.835 | 0.778 | 0.898 |

Analysing the impact of self-attention on a per-species level shows a consistent positive effect of the inclusion of self-attention across all species (Fig. 4A). The improvement in average precision is particularly noticeable for species that exhibited lower performance without the self-attention. For instance, the ability of the model to classify *Myotis* species, which encompasses several challenging and previously poorly-performing species, was significantly improved. In contrast, removing the *CoordConv* layers resulted in a negligible impact on overall performance across all species, although a slight improvement was observed for the two lowest-performing *Myotis* species (Fig. 4B).

**FIGURE 4.**
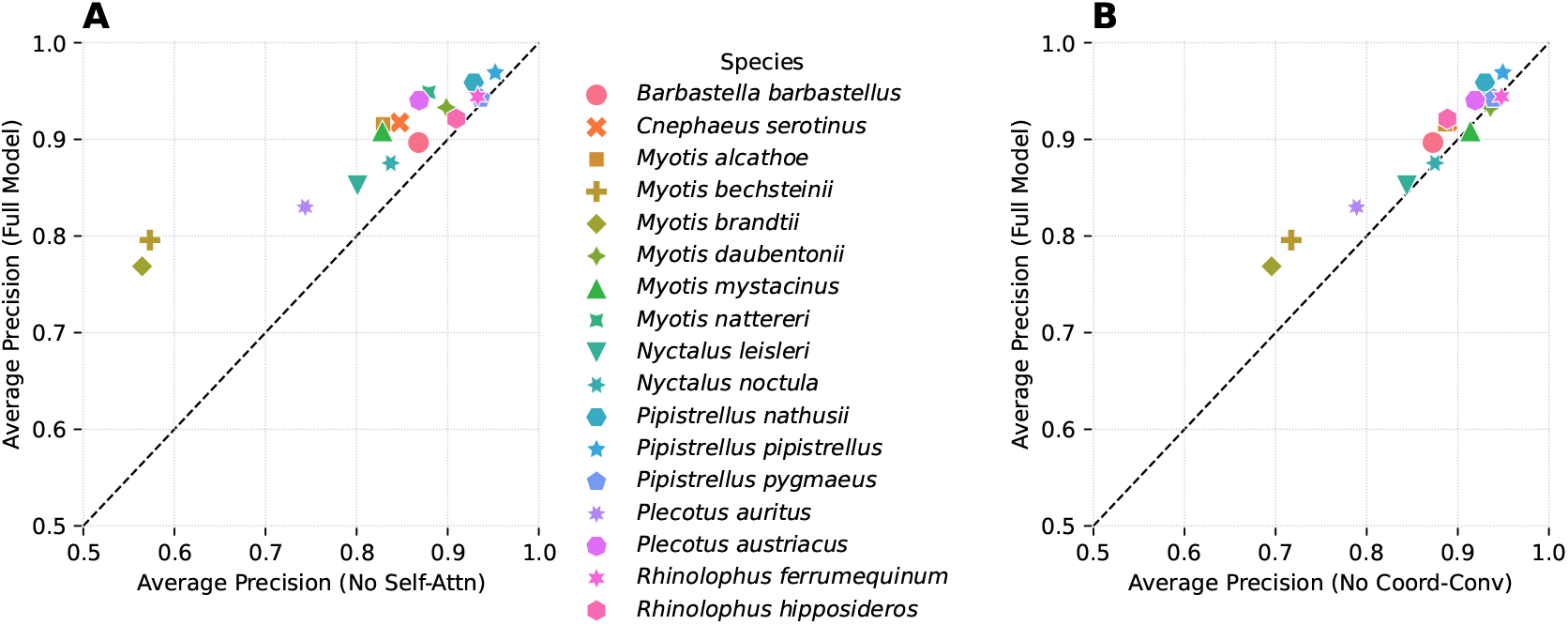
Impact of BatDetect2 modifications on per-species average precision. (A) Average precision (AP) for each species in the UK_same_ dataset for the full BatDetect2 model and the NoSelfAttn variant (without self-attention). Points above the diagonal line indicate superior performance of the full model. (B) AP for each species in the UK_same_ dataset for the full BatDetect2 model and the NoCoordConv variant (without frequency information added to the convolutional layers). All models were trained three times on the UK_same_ training split, and the mean AP on the corresponding test split is reported.

### 3.3 Detection performance in novel regions

We found that BatDetect2_UK_, the model trained solely on the UK_same_ dataset and applied directly to the target datasets, exhibited strong detection performance in all but one case (Table 3). In the Australia dataset, BatDetect2_UK_ showed strong detection performance (AP Det = 0.982), achieving a score comparable to the BatDetect2 model trained directly on Australian data. Performance on the Yucatan dataset was also encouraging (AP Det = 0.821). In contrast, performance declined markedly in the Brazil dataset, where BatDetect2_UK_ performance was significantly lower (AP Det = 0.573). Despite this reduced accuracy, BatDetect2_UK_ still outperformed the Random Forest (RF) baseline trained on the same UK_same_ dataset in terms of AP Det.

**TABLE 3.**
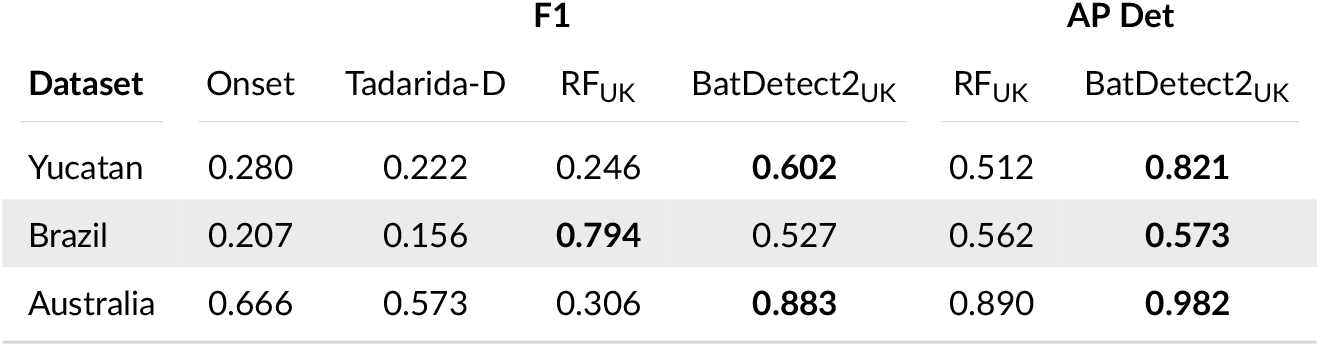
Evaluation of BatDetect2 detection performance in three novel regions. Models are evaluated using Average Precision (AP Det) and F1 score. The BatDetect2_UK_ and RF_UK_ models were trained solely on UK_same_ and applied to the target datasets without further training. Onset is a simple onset detector based on the strength envelope, and Tadarida-D is used out of the box without any adjustment. Note that AP Det is not computed for the Onset and Tadarida-D methods as they do not provide a confidence score. For F1 score computation, an illustrative threshold of 0.5 was chosen.

With respect to F1-score, BatDetect2_UK_ substantially exceeded the two generic detection approaches, Onset and Tadarida-D, though RF_UK_ achieved a higher F1-score in the Brazil dataset. Note that we used the default confidence threshold of 0.5 and did not optimise thresholds for individual models. Therefore, we consider AP Det to provide a more reliable, threshold-independent measure of performance. For both the Yucatan and Brazil datasets, the model trained directly on the respective regional data consistently outperformed BatDetect2_UK_ and all other evaluated approaches.

## 4 DISCUSSION

We have shown that the full BatDetect2 model is able to learn to detect and classify echolocation calls from bats across five different datasets. BatDetect2 significantly outperforms the traditional call parameter-based baseline, providing a strong argument in favour of DL models over traditional methods for this task. Despite the growing trend towards DL methods, direct comparisons with call parameter-based methods remain scarce, for example only in Mac Aodha et al. (2018) for detection and Fundel et al. (2023) for classification.

For the majority of species in the UK_same_ dataset split, BatDetect2 results in high precision at high recall rates (Fig. 3). This is important as it enables practitioners to trade-off recall for precision to ensure that they obtain reliable, high confidence, predictions from the model. The file-level mean average precision is 81.6% and 91.4% for the UK_diff_ and UK_same_ datasets, where a large percentage of the mistakes can be attributed to known challenging species, i.e., the *Myotis* species (Parsons and Jones, 2000; Walters et al., 2012). Although no clear relationship between the number of training examples and performance was observed for the UK_same_ dataset, the higher performance of the model trained on UK_same_ suggests that larger and more representative training datasets can improve model robustness. However, it is difficult to disentangle the effect of increased training data size from the more rigorous, independent train-test split used for UK_diff_, where recordings from the same source were not shared between training and testing sets. Further investigation is warranted to clarify these effects.

We showed that by modifying the base CNN architecture with the addition of a self-attention layer considerably improves classification performance. This improvement is particularly pronounced for species within the *Myotis* genus, suggesting that leveraging the temporal structure of call sequences may be crucial for accurate classification within this group. Through its self-attention layer, BatDetect2 efficiently utilises information from longer input timescales without significantly increasing computational cost, resulting in a model that can perform inference approximately 17 times faster than real-time on a GPU. However, modifications aimed at enhancing the use of frequency data did not yield notable performance improvements. Although the proposed modification was relatively simple, and other approaches might enhance performance, it is also possible that a fully CNN model already effectively captures this frequency information. This aligns with findings that CNNs, while translation invariant in theory, often exhibit subtle violations of this property in practice (Zhang, 2019). Overall, these findings highlight the importance of incorporating longer audio context and long-range temporal patterns into the design of deep learning models for bat call classification, and potentially for other bioacoustic tasks. Further exploration of audio-specific architectures, particularly those using raw audio directly instead of 2D image representations (Ravanelli and Bengio, 2018; Hagiwara, 2023), holds significant potential for further performance improvements (Stowell, 2022).

While the pre-trained BatDetect2 model displayed varied performance across novel regions, its general detection capability remains promising, as evidenced by high detection metrics (AP Det) in the Yucatan and Australia datasets. However, performance on the Brazil dataset was significantly lower, potentially due to the abundance of low-frequency calls in that region (Fig. S6). Since the UK training dataset is primarily composed of echolocation calls above 25 kHz (Fig. S3), this could have introduced a bias, causing the model to under-detect calls below this frequency threshold when applied to the Brazilian data. Despite this limitation, BatDetect2 outperformed all other tested solutions, representing a valuable improvement for practitioners detecting individual echolocation calls. Furthermore, this shows promise that including more data for pre-training could lead to a robust general echolocation detection model.

BatDetect2 performs well across the five datasets tested, however it still relies on the availability of diverse, and exhaustively annotated, training data. Collecting such audio data can be challenging, in addition to being time-consuming to annotate. While methods for semi-supervised and self-supervised training offer the potential to learn effective models with limited to no training supervision (Heggan et al., 2024; Hagiwara et al., 2023), diverse labelled data is still needed to evaluate the performance of the developed models. Bat calls can exhibit plasticity depending on the population sampled (Montauban et al., 2021), the presence of other species, and the composition of the local environment. As a result, care needs to be taken to ensure that the collected training datasets are representative of the downstream deployment situations as much as possible (van Merriënboer et al., 2024). Finally, our training datasets currently only contain annotated echolocation calls, and thus the model cannot make predictions for other types of calls, e.g., social calls or feeding buzzes. This limitation could be addressed with appropriate training data.

Unlike typical deep learning-based classifiers, BatDetect2 returns a list of detections for a given input recording, each represented by a time-frequency bounding box around the detected call and associated species scores (Fig. 2). This is valuable as it enables easier inspection of the model’s predictions, facilitating a better understanding of potential failure cases. However, it is left up to the user to decide how to best merge the individual detections into a set of ‘bat passes’, where a pass constitutes a sequence of individual calls. This aggregation step is often crucial, as downstream analysis are typically derived from the number of detected individuals or their activity levels (e.g., Ferreira et al. (2022) and Hoggatt et al. (2024)), rather than the number of detected calls. One approach is to use a grouping-based heuristic based on the time between detected calls as in Mac Aodha et al. (2018). The high recall rates of BatDetect2 means that this type approach is less likely to separate individual bat passes into multiple different ones. In contrast, methods that produce high numbers of false negatives run the risk of over-counting the number of passes as they can miss faint calls in a sequence, and thus incorrectly break them up into a number of shorter passes. Still, a better understanding of how to best merge these detections into passes is needed, particularly for distinguishing calls from individual bats. This would allow for more accurate estimates of the number of individuals present, improving the reliability of population monitoring.

In this study we demonstrate that the proposed training pipeline can be applied to audio data from distinct regions without requiring modifications to the underlying code. This pipeline is packaged in the open-source Python package `batdetect2`, available on GitHub. In conjunction with accessible annotation tools like `whombat` (Martínez Balvanera et al., 2024), `batdetect2` enables the training and deployment of models on custom annotated datasets, even beyond bat species. However, while no coding is required to train a model, some technical expertise is still needed to set up the training environment and to understand the model’s output. Integrating the training pipeline into user-friendly, graphical interface tools could offer a more accessible solution for practitioners. Ultimately, this work helps to democratise the development of specialised bioacoustic models by removing significant technical barriers, thereby enabling practitioners to focus on collecting and annotating datasets for their species of interest. Coupled with efficient annotation workflows (Eichinski et al., 2022; Dumoulin et al., 2025), our pipeline will help accelerate the broader development of targeted bioacoustic models.

## Acknowledgements

We thank Giada Giacomini, Annika Binet, Michelle Parsons, and Steve Lucas for their invaluable assistance with the annotation efforts for the UK datasets. We also thank Stuart Newson for collecting and sharing recordings from the UK with the Bat Detective project, and for permitting their use in this study. For the Brazilian data, we thank Liz Walsh for assistance with annotation, and Adriana Arias-Aguilar for help with the genus identification of Brazilian bat calls. Collection of bat calls in Australia was supported by funding from the National Landcare Program and the Cotton Research and Development Corporation. This work was partially supported by Secretaría de Ciencia, Humanidades, Tecnología e Innovación (SECIHTI) through a PhD scholarship (Grant/Award Number: 2020-000017-02EXTF-00334). We are also grateful to Grant Van Horn for helpful discussions.

## Author Contribution

**Conceptualization**: OMA, SMB, KEJ;

**Data curation**: OMA, SMB, ED, MC, PE, EB, AS, MB, RC, GG, MCMG, MKO, SP, TS;

**Formal analysis**: OMA, SMB;

**Funding acquisition**: OMA, LG, KB, KEJ;

**Investigation**: OMA, SMB, ED, MC, MB, RC, MCMG, MKO, SP, TS;

**Methodology**: OMA, SMB, KEJ;

**Project administration**: OMA, LG, KB, GG, KEJ;

**Resources**: OMA, LG, KB, KEJ;

**Software**: OMA, SMB, PE;

**Supervision**: OMA, KEJ;

**Visualization**: OMA, SMB;

**Writing – original draft**: OMA, SMB, KEJ;

**Writing – review & editing**: OMA, SMB, ED, MC, PE, EB, LG, AS, MB, KB, RC, GG, MCMG, MKO, SP, TS, KEJ.

**OMA**: Oisin Mac Aodha **SMB**: Santiago Martínez Balvanera **ED**: Elise Damstra **MC**: Martyn Cooke **PE**: Philip Eichinski **EB**: Ella Browning **LG**: Lia Gilmour **AS**: Agata Staniewicz **MB**: Michel Barataud **KB**: Katherine Boughey **RC**: Roger Coles **GG**: Giada Giacomini **MCMG**: M. Cristina Mac Swiney G. **MKO**: Martin K. Obrist **SP**: Stuart Parsons **TS**: Thomas Sattler **KEJ**: Kate E. Jones

## Supporting Information

### A PERFORMANCE REPORT

In this section, we present a comprehensive report of the BatDetect2 performance across all five datasets (Fig. S1). For each dataset, precision-recall (PR) curves are presented for each species. Additionally, per-genus PR curves are displayed. Finally, file-level confusion matrices are presented.

To produce the genus-level PR curves, both the ground truth annotations and the model predictions must be adapted. First, all ground truth species labels are converted to their corresponding genus level. Next, we generate a genus-level confidence score for each prediction. This is achieved by summing the per-species confidence scores corresponding to all species that belong to the given genus. The computation of the PR curves then follows the standard procedure described previously for mAP Class. Specifically, for each genus, we match ground truth calls and predictions based on temporal overlap. Priority for matching is determined by the genus-level confidence score.

The confusion matrices are generated by matching predictions and ground truth annotations and then comparing the true species against the species with the highest predicted confidence (top confidence). The priority for matching a prediction is determined by the top confidence score. We exclude non-matched predictions or ground truth annotations (i.e., false positives and false negatives), as well as any matches involving annotations where the species identity is unknown. Furthermore, predictions with a top confidence score below 0.2 are also excluded. Finally, the rows are normalised to sum to 1. Consequently, a cell with value *x* at the intersection of row (Species 1) and column (Species 2) is interpreted as: “Of all true calls belonging to Species 1, a proportion of *x* were predicted as Species 2.”

**FIGURE S1.**
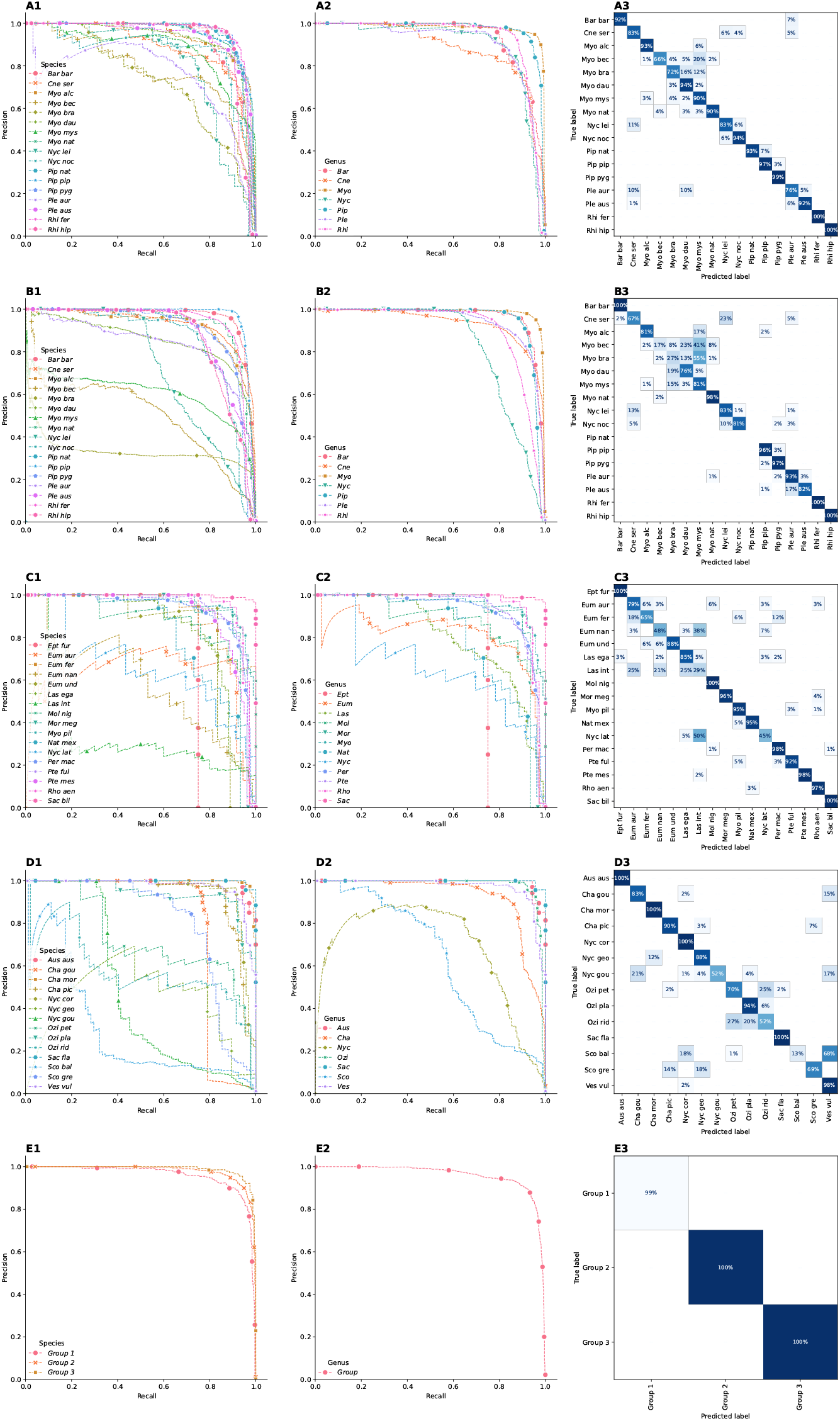
Precision-recall (PR) and confusion matrices for our BatDetect2 model for the five different test sets: UK_same_ (A1-3), UK_diff_ (B1-3), Yucatan (C1-3), Australia (D1-3) and Brazil (E1-3) respectively. The first column (1) depicts the per-species precision-recall curves and the second column (2) is the per-genus equivalent. The third column (3) illustrates the call-level confusion matrix, where white rows indicate that there were no species of that type in the filtered test set.

### B ATTENTION MECHANISM

This section provides an illustrative example of the self-attention mechanism employed within the BatDetect2 model. During the processing of an input spectrogram, the self-attention mechanism enables the model to identify and weight the most relevant time steps for predicting the species present at a given time point. The attention module operates in the bottleneck of the network, where the spectrogram has been downsampled and transformed into a temporal sequence of 256-dimensional feature vectors, each representing a time step of approximately 8 ms. It operates solely along the temporal dimension. For each time step, the module computes self-attention scores against all other time steps in the input sequence. This process allows the model to leverage global contextual information across the entire input sequence when estimating the species present at a specific time point (Fig. S2).

**FIGURE S2.**
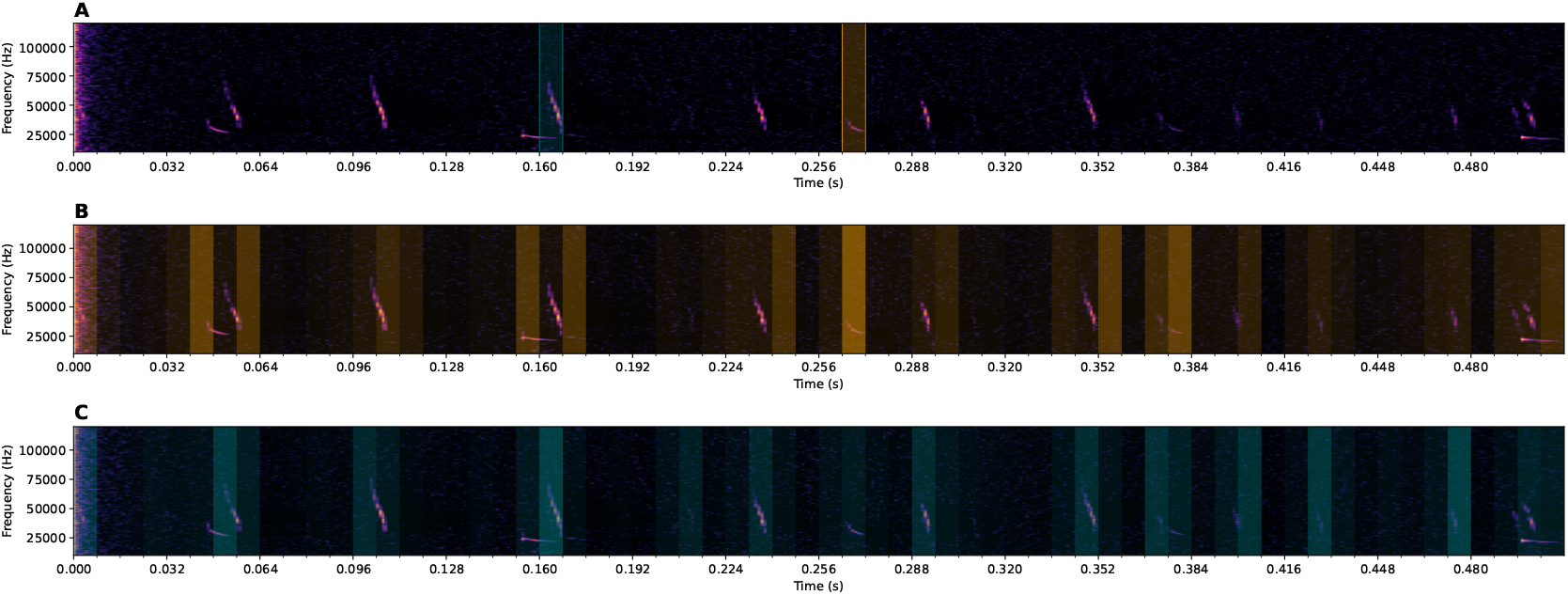
Visualisation of the self-attention scores. (A) Example input spectrogram from the UK_same_ training set, with two query locations highlighted in orange and teal. (B, C) The same spectrogram with temporal regions shaded according to the attention weights assigned when the model attends to the orange query (B) and the teal query (C). Greater colour intensity indicates that the model places more weight on that temporal region when forming its prediction for the corresponding query location. The teal query contains a *Myotis daubentonii* call, and other calls from this species receive high attention. A similar pattern is observed for the *Nyctalus leisleri* call in the orange query region, although additional portions of the spectrogram are also highlighted. Note that the highlighted temporal regions (including the query points) have lower temporal resolution than the spectrogram because they correspond to the downsampled bottleneck features, with a resolution of 8 ms per region.

### C AUDIO DATASETS

Here we provide additional details of the different datasets used in for training and evaluation.

#### C.1 UK data

As mentioned in the main text the UK datasets is composed of recordings with calls of known species, recordings with calls from unknown species and empty recordings. In total there are 17 species in the UK dataset (Fig. S3). This is the total number of species which are known to be breeding in the UK.

The annotation process prioritised annotating only one clip, at most two seconds in duration, from each original input recording, rather than densely annotating long, multi-second audio files. This was also performed in order to increase the data diversity, as there can often be a large amount of self-similarity within the same longer recording. As a result, the clipped files vary in duration from between 0.4 to two seconds, and the average duration is just over one second.

The BatDetect2 model predicts the location of the lower left corner for each echolocation call in an input recording. For the two constant call frequency-based species in the UK, *Rhinolophus ferrumequinum* and *Rhinolophus hipposideros*, there was a high degree of variability in the position of the lower left corner of the call. This happens as a direct result of the recording quality, characteristics of the local environment, and the distance of the bat from the microphone As a result, it was often difficult to determine the exact lower frequency for these two species. To overcome this issue, we standardised the lower and upper frequency for each of the these species by setting them to per-species mean values, where the means were computed on the training sets.

We constructed two splits for the UK dataset. Both splits contain the same number of calls overall, and only differ in how the data is distributed between their respective training and test sets. As noted earlier, there are six main sources of data for the UK bat recordings. The first split, referred to as UK_same_, simply shuffles the files randomly into training and test sets and ensures that there is a maximum of four recordings (i.e. files not calls) per species, per data source, in the test set. This results in a split with 7,010 training files and 369 test files (Table S1).

The second split, UK_diff_, is more challenging. Here we simulate a difficult real world setting where an entire data source is held out for validation. We remove one of the largest sources, which leaves 5,911 training files and 1,468 test files (Table S2). This increases the difficulty due to the reduction in the training set size as well as increasing any potential domain gap that may exists between the train and test sets. This test set does not contain one of the species, *Pipistrellus nathusii*, as it was not possible to capture any recordings of it. Note that in both cases the data is still split at the file level (as opposed to individual call level). This minimises any potential overlap between the training and test sets.

#### C.2 Yucatan data

This dataset consists of 285 passive recordings gathered in the Yucatan peninsula in Mexico as part of a field study conducted between 2004 and 2006 (MacSwiney G. et al., 2008). A Pettersson D980 bat detector device was used to detect and record bat calls. The device was active throughout three ten-minute periods at night, in a total of eight sites and covering twelve sampling nights per site. When active, and if a bat call was detected, the device would record for three seconds and a time expanded version would be stored on a magnetic tape. The recordings were then cut into one second clips, resulting in a total of 1,193 audio files.

The species identification of the bat calls was made in two phases. For the original study, all recordings were reviewed manually. From each recording, at most five representative echolocation calls per detected species was selected and analyzed using Bat Sound Pro 3.10. The species of each call was then identified through comparison to a bat call library of captured bats from the same study. Please consult MacSwiney G. et al., 2008 to see the full details of their identification protocol.

**FIGURE S3.**
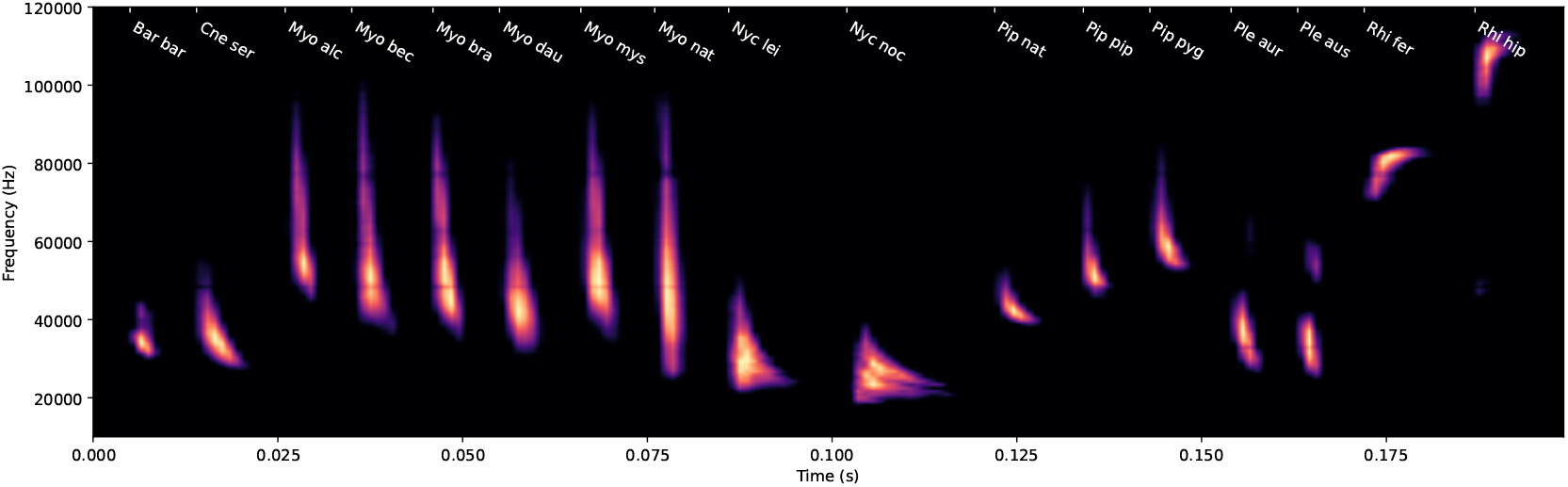
Average echolocation calls for *UK*_same_ species. This figure visualises the average spectrogram for each species from the *UK*_same_ training dataset. These averages were computed by aligning the spectrogram of each individual call and then averaging them. All spectrograms were generated using the same preprocessing steps as the input to BatDetect2. The vertical axis represents frequency (10–120 kHz) and the horizontal axis represents time in seconds. Note that this averaging process can hide the substantial intra-species call variation. For instance, the constant frequency (CF) component and subsequent downward sweep of the two *Rhinolophus* species disappear due to the high variability in the CF section’s duration.

In the second phase we annotated all missing bat calls using the annotation interface. Bounding boxes were drawn around each detected bat call in the spectrogram. Species identification was performed by comparing to the previously annotated calls. In cases where it was not possible to determine the species, the call was labelled using the generic ‘Bat’ class. A recording was fully annotated when all bat echolocation calls were marked with a bounding box and all recognisable calls were tagged with its species, or the generic, label. This resulted in a total of 1,193 audio clips that were fully annotated and kept as part of the dataset. Three species (*Pteronotus personatus, Molossops greenhalli*, and *Molossus sinaloae*) were excluded as they only appeared in fewer than seven distinct recordings. The annotations for these species was set to the generic ‘Bat’ class. The final annotated dataset consists of 10,020 individual bat echolocation calls with bounding box annotations from 17 different species (Fig. S4).

To train and evaluate the detection and classification models we split the dataset into distinct training and testing subsets. To minimise any leakage from the test to the train set, we opted to split the data at the recording level, i.e. we avoided including one-second clips from the same recording in the training and testing subsets. The test set contains ~20% (282 audio clips) of all recordings while the remaining ~80% (911 audio clips) was used for training (Table S3). In order to maintain the distribution of calls per species between the full dataset and the testing and training datasets, we labelled each recording with all its occurring species and used a stratified sampling method for multilabel datasets (Sechidis et al., 2011).

#### C.3 Australia data

The Australian dataset used to train and test the model was taken from a bat call reference library collected by a bat expert. The subset used consists of a set of 14 bat species which have a sympatric distribution in the major cotton growing region on the north west plains of New South Wales and adjacent areas in central southern Queensland. Bat calls were recorded in the field from individuals released after capture, following positive species identification. A custom made digital ultrasound recorder from Nanobat Systems was used to record echolocation calls in 5 second sequences with a sampling rate 500 kHz and stored as 16 bit WAVs. Bats were recorded for as long as they flew around the release site until out of recording range. The resulting files were analysed and edited using Audacity 3.2.0 to find echolocation pulse sequences with good signal to noise ratio, undistorted waveforms and as close to search phase as possible. Edited wav files were then accumulated from the release recordings of multiple individuals of the same species and across the species group.

**FIGURE S4.**
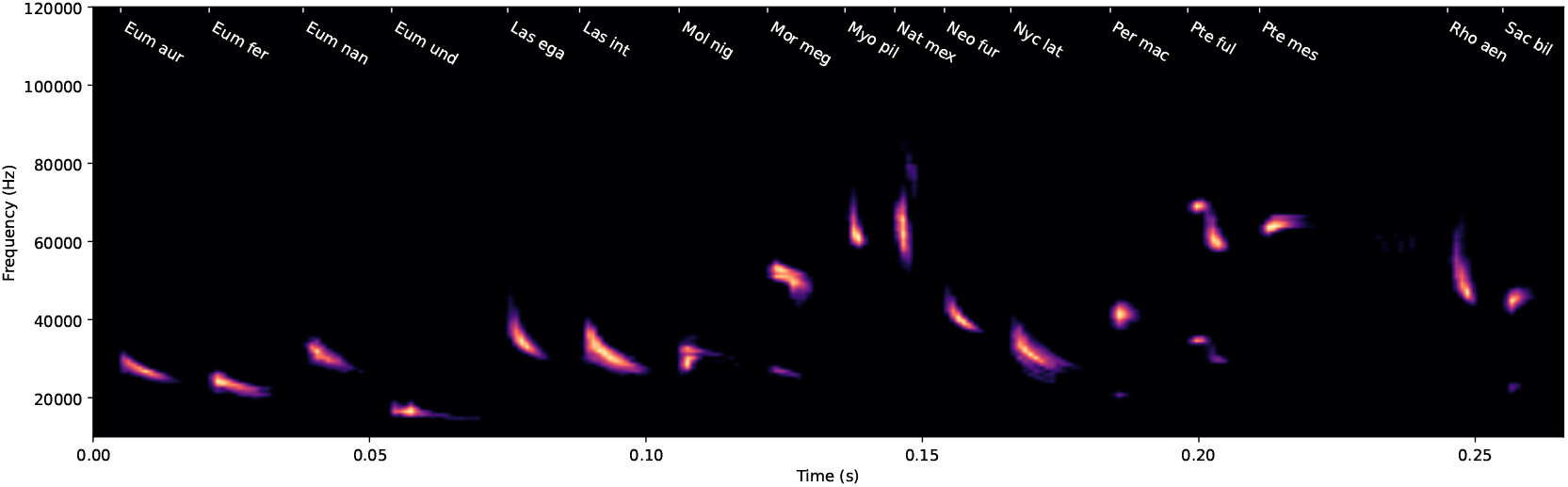
Average echolocation calls for Yucatan species. This figure visualises the average spectrogram for each species from the Yucatan training dataset. These averages were computed by aligning the spectrogram of each individual call and then averaging them. All spectrograms were generated using the same preprocessing steps as the input to BatDetect2. The vertical axis represents frequency (10–120 kHz) and the horizontal axis represents time in seconds. Note that this averaging process can hide the substantial intra-species call variation.

These audio files had an average length of 3.29 seconds, with the shortest being 0.23 seconds and the longest being 10 seconds in duration. All annotated pulses were labelled by species since the original sequences were obtained from individually released bats, identified to species level. The only exception comes from the Ozimops species where the low release number of individuals (rarely caught) was augmented by identifying species from additional field recordings of bat activity at night. This was done manually by conventional sound analysis of field recordings taken from various study areas and using an experienced bat bioacoustics expert familiar with this genus. There were some instances where multiple species may have been present in a given file, and thus were potentially incorrectly attributed to the wrong species label.

The data was randomly split at the file level, with 80% of the recordings for a species staying the train set, and the rest in the test. This resulted in 220 training and 60 testing files (Table S4).

#### C.4 Brazil data

Data for this study was collected between January and March 2019 in south-eastern Brazil. The data used for training is a subset of acoustic data collected using AudioMoth (Hill et al., 2019) recorders which were set to record at a sampling rate of 395 kHz for one minute every five minutes between 22:00 and 04:00. The recorders were deployed on coffee plantations and in adjacent forest fragments. The final dataset consists of 320 ten second audio recordings. As no species labels were available for this dataset, we opted to group the calls into groups based on their dominant frequency. Calls were initially labelled at the family or genus level, where recording quality allowed, and were later merged into coarser call-type groups. Genus-level identification was conducted with the help of experts with extensive experience in Brazilian bat call identification (Arias-Aguilar et al., 2018). This resulted in three distinct sonotypes (Fig. S6), along with the generic bat class which served as an additional class for cases where it was not possible to identify calls to one of the previously mentioned three groups.

**FIGURE S5.**
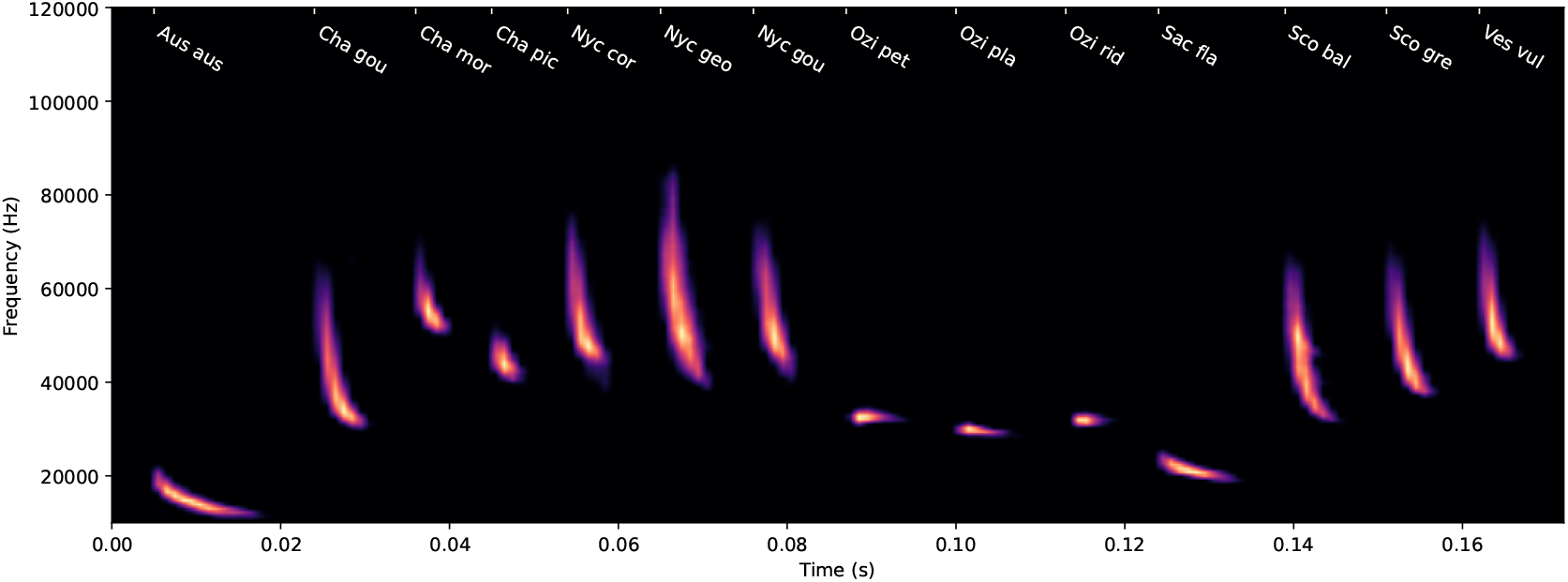
Average echolocation calls for Australian species. This figure visualises the average spectrogram for each species from the Australian training dataset. These averages were computed by aligning the spectrogram of each individual call and then averaging them. All spectrograms were generated using the same preprocessing steps as the input to BatDetect2. The vertical axis represents frequency (10–120 kHz) and the horizontal axis represents time in seconds. Note that this averaging process can hide the substantial intra-species call variation.

We randomly assigned ~80% of the audio files (256 files) to the training set and the remaining ~20% (64 files) to the test set. This resulted in a total of 7,989 and 2,010 calls in the respective sets (Table S5).

**FIGURE S6.**
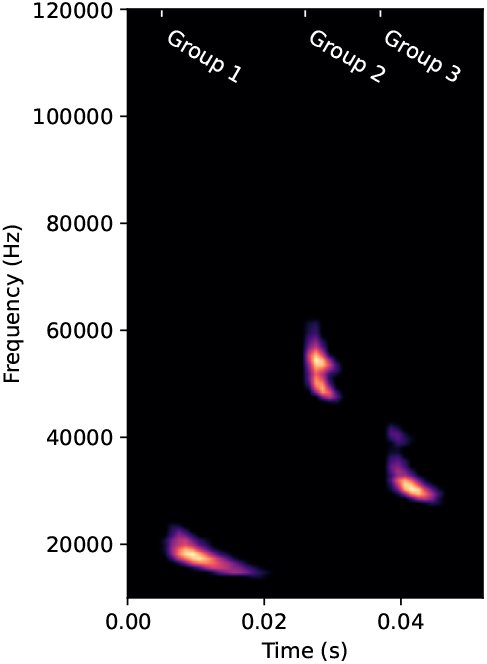
Visualisation of the Brazil data. This figure visualises the average spectrogram for each species group from the Brazil training dataset. These averages were computed by aligning the spectrogram of each individual call and then averaging them. All spectrograms were generated using the same preprocessing steps as the input to BatDetect2. The vertical axis represents frequency (10–120 kHz) and the horizontal axis represents time in seconds. Note that this averaging process can hide the substantial intra-species call variation.

**TABLE S1.**
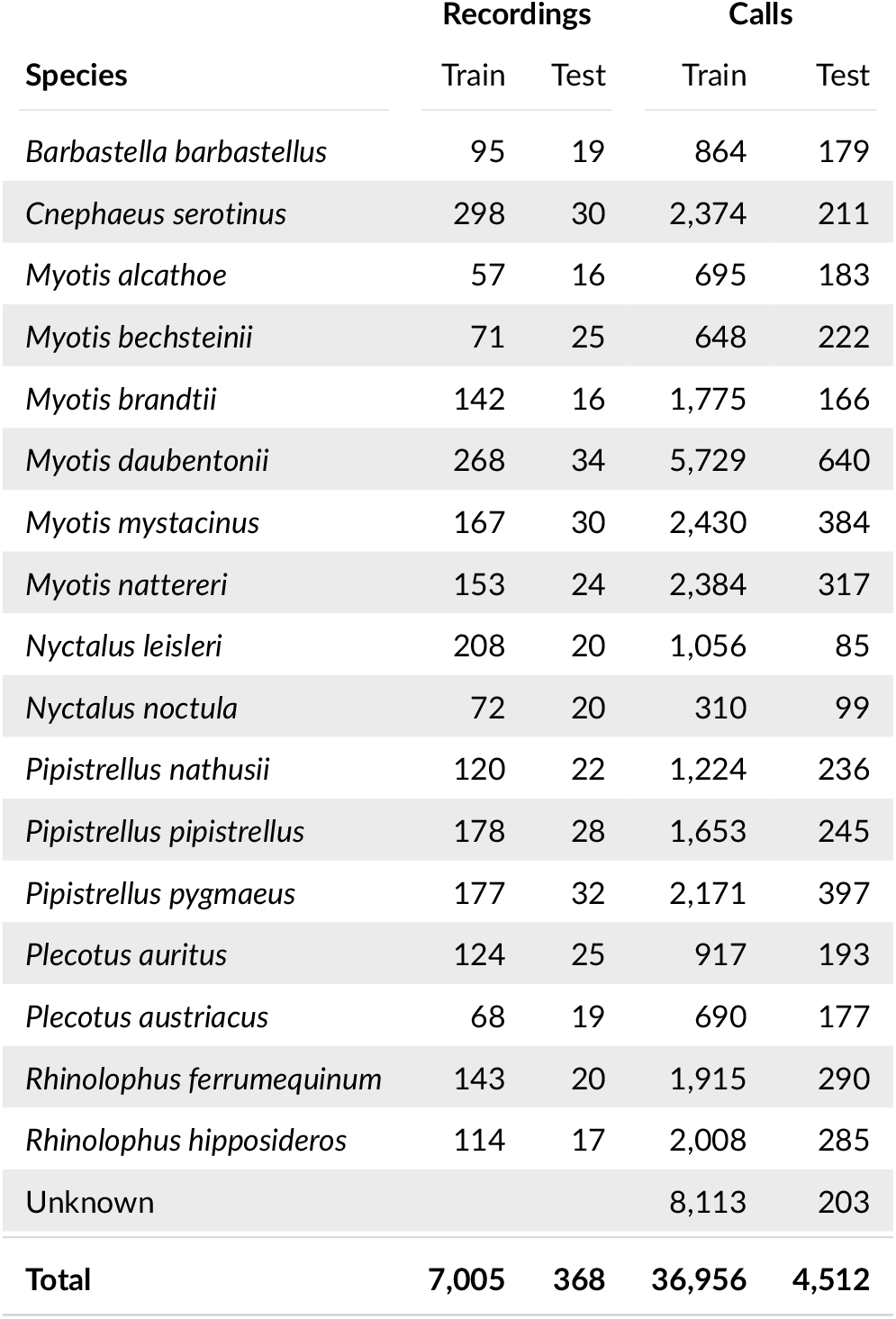
Summary of the UK_same_ Dataset. The Recordings columns show the number of recordings in which a call of the given species appears. Note that a single recording may contain multiple species, so the sum may exceed the total recording count. The Calls columns show the number of individual echolocation calls annotated for each species. This includes calls for which a species could not be assigned with certainty (Unknown). The totals for recordings and annotated calls across the entire test and train sets are provided in the final row.

**TABLE S2.** Summary of the UK_diff_ Dataset. The Recordings columns show the number of recordings in which a call of the given species appears. Note that a single recording may contain multiple species, so the sum may exceed the total recording count. The Calls columns show the number of individual echolocation calls annotated for each species. This includes calls for which a species could not be assigned with certainty (Unknown). The totals for recordings and annotated calls across the entire test and train sets are provided in the final row.

| Species | Recordings |  | Calls |  |
| --- | --- | --- | --- | --- |
|  | Train | Test | Train | Test |
| *Barbastella barbastellus* | 57 | 57 | 468 | 575 |
| *Cnephaeus serotinus* | 80 | 248 | 403 | 2,182 |
| *Myotis alcathoe* | 29 | 44 | 374 | 504 |
| *Myotis bechsteinii* | 22 | 74 | 241 | 629 |
| *Myotis brandtii* | 30 | 128 | 351 | 1,590 |
| *Myotis daubentonii* | 156 | 146 | 3,998 | 2,371 |
| *Myotis mystacinus* | 82 | 115 | 1,378 | 1,436 |
| *Myotis nattereri* | 169 | 8 | 2,610 | 91 |
| *Nyctalus leisleri* | 117 | 111 | 695 | 446 |
| *Nyctalus noctula* | 66 | 26 | 209 | 200 |
| *Pipistrellus nathusii* | 142 | 0 | 1,460 | 0 |
| *Pipistrellus pipistrellus* | 117 | 89 | 868 | 1,030 |
| *Pipistrellus pygmaeus* | 155 | 54 | 1,461 | 1,107 |
| *Plecotus auritus* | 82 | 67 | 528 | 582 |
| *Plecotus austriacus* | 32 | 55 | 331 | 536 |
| *Rhinolophus ferrumequinum* | 42 | 121 | 717 | 1,488 |
| *Rhinolophus hipposideros* | 40 | 91 | 722 | 1,571 |
| Unknown |  |  | 7,492 | 814 |
| **Total** | **5,906** | **1,467** | **24,306** | **17,152** |

**TABLE S3.** Summary of the Yucatan Dataset. The Recordings columns show the number of recordings in which a call of the given species appears. Note that a single recording may contain multiple species, so the sum may exceed the total recording count. The Calls columns show the number of individual echolocation calls annotated for each species. This includes calls for which a species could not be assigned with certainty (Unknown). The totals for recordings and annotated calls across the entire test and train sets are provided in the final row.

| Species | Recordings |  | Calls |  |
| --- | --- | --- | --- | --- |
|  | Train | Test | Train | Test |
| *Eumops auripendulus* | 59 | 16 | 156 | 36 |
| *Eumops ferox* | 37 | 13 | 60 | 24 |
| *Eumops nanus* | 23 | 9 | 66 | 33 |
| *Eumops underwoodi* | 28 | 10 | 36 | 18 |
| *Lasiurus ega* | 42 | 15 | 250 | 69 |
| *Lasiurus intermedius* | 28 | 10 | 106 | 31 |
| *Molossus nigricans* | 15 | 6 | 65 | 25 |
| *Mormoops megalophylla* | 28 | 6 | 172 | 30 |
| *Myotis pilosatibialis* | 52 | 12 | 519 | 90 |
| *Natalus mexicanus* | 9 | 5 | 62 | 26 |
| *Neoptesicus furinalis* | 22 | 1 | 94 | 4 |
| *Nyctinomops laticaudatus* | 33 | 7 | 98 | 23 |
| *Peropteryx macrotis* | 215 | 69 | 1,036 | 322 |
| *Pteronotus fulvus* | 64 | 18 | 509 | 166 |
| *Pteronotus mesoamericanus* | 38 | 9 | 345 | 81 |
| *Rhogeessa aenea* | 20 | 4 | 166 | 36 |
| *Saccopteryx bilineata* | 49 | 11 | 381 | 92 |
| Unknown | 405 | 122 | 3,710 | 1,291 |
| Total | 911 | 282 | 7,831 | 2,397 |

**TABLE S4.** Summary of the Australian Dataset. The Recordings columns show the number of recordings in which a call of the given species appears. Note that a single recording may contain multiple species, so the sum may exceed the total recording count. The Calls columns show the number of individual echolocation calls annotated for each species. This includes calls for which a species could not be assigned with certainty (Unknown). The totals for recordings and annotated calls across the entire test and train sets are provided in the final row.

| Species | Recordings |  | Calls |  |
| --- | --- | --- | --- | --- |
|  | Train | Test | Train | Test |
| *Austronomus australis* | 8 | 2 | 125 | 35 |
| *Chalinolobus gouldii* | 37 | 10 | 568 | 138 |
| *Chalinolobus morio* | 16 | 4 | 429 | 117 |
| *Chalinolobus picatus* | 13 | 4 | 327 | 92 |
| *Nyctophilus corbeni* | 17 | 4 | 537 | 95 |
| *Nyctophilus geoffroyi* | 8 | 1 | 179 | 19 |
| *Nyctophilus gouldi* | 20 | 6 | 363 | 105 |
| *Ozimops petersi* | 8 | 2 | 149 | 42 |
| *Ozimops planiceps* | 10 | 3 | 142 | 51 |
| *Ozimops ridei* | 8 | 3 | 122 | 48 |
| *Saccolaimus flaviventris* | 9 | 3 | 131 | 23 |
| *Scotorepens balstoni* | 8 | 2 | 232 | 82 |
| *Scotorepens greyii* | 13 | 4 | 273 | 76 |
| *Vespadelus vulturnus* | 35 | 9 | 812 | 121 |
| Unknown |  |  | 180 | 111 |
| Total | 216 | 58 | 4,569 | 1,154 |

**TABLE S5.** Summary of the Brazil Dataset. The Recordings columns show the number of recordings in which a call of the given call type appears. Note that a single recording may contain multiple call types, so the sum may exceed the total recording count. The Calls columns show the number of individual echolocation calls annotated for each identified call type. This includes calls for which a type could not be assigned (Unknown). The totals for recordings and annotated calls across the entire test and train sets are provided in the final row.

| Species | Recordings |  | Calls |  |
| --- | --- | --- | --- | --- |
|  | Train | Test | Train | Test |
| Group One | 122 | 32 | 2,168 | 490 |
| Group Two | 74 | 18 | 2,992 | 742 |
| Group Three | 43 | 6 | 1,182 | 159 |
| Unknown |  |  | 1,646 | 619 |
| Total | 256 | 64 | 7,989 | 2,010 |

**TABLE S6.** Description of the full architecture for BatDetect2 model. Input and output sizes are (feature dimension, height, width), where height and width correspond to frequency and time bins in the spectrogram, respectively. Although the model has no strict requirement for input width, we use 512 for illustration, which corresponds to 512ms of audio given the preprocessing pipeline. The kernel size is represented as (height, width). In the case where two tensors are added together for the input to a layer, this is simply performed using an element wise addition. The model outputs a *C* + 1 dimensional vector for each location in time and frequency, where *C* + 1 represents the number of classes plus one additional class for background, i.e. ‘Not bat’. The model also outputs an additional two dimensional vector for each location which encodes the predicted width (i.e. duration) and height (i.e. frequency range) of any echolocation event at that location in time and frequency. The parameter count for most layers is independent of the number of classes (C). Only the final classification layer (*Ŷ*) depends on C. The counts shown are illustrative, based on C=17 UK species; the exact formula is provided in brackets. The total parameter count therefore depends on C, requiring only small adjustments if a different number of classes is used.

| Layer Name | Input | Layer Type | Input Size | Output Size | Kernel Size | Num Parameters |
| --- | --- | --- | --- | --- | --- | --- |
| **Encoder** |  |  |  |  |  |  |
| conv_down_0 | spectrogram | CoordConvDown | (1, 128, 512) | (32, 64, 256) | (3,3) | 800 |
| conv_down_1 | conv_down_0 | CoordConvDown | (32, 64, 256) | (64, 32, 128) | (3,3) | 19,264 |
| conv_down_2 | conv_down_1 | CoordConvDown | (64, 32, 128) | (128, 16, 64) | (3,3) | 75,296 |
| conv_3 | conv_down_2 | Conv2d, BN, ReLU | (128, 16, 64) | (256, 16, 64) | (3,3) | 295,680 |
| **Bottleneck** |  |  |  |  |  |  |
| conv_1d | conv_3 | Conv2d, BN, ReLU | (256, 16, 64) | (256, 1, 64) | (16,1) | 1,049,344 |
| self_attn | conv_1d | Self-Attention | (256, 1, 64) | (256, 1, 64) | n/a | 263,168 |
| repeat_vert | self_attn | Repeat Vertical | (256, 1, 64) | (256, 16, 64) | n/a | 0 |
| **Decoder** |  |  |  |  |  |  |
| conv_up_0 | repeat_vert + conv_3 | CoordConvUp | (256, 16, 64) | (64, 32, 128) | (2,2) | 148,256 |
| conv_up_1 | conv_up_0 + conv_down_1 | CoordConvUp | (64, 32, 128) | (32, 64, 256) | (2,2) | 18,880 |
| conv_up_2 | conv_up_1 + conv_down_0 | CoordConvUp | (32, 64, 256) | (32, 128, 512) | (2,2) | 9,728 |
| conv_op_0 | conv_up_2 | Conv2d, BN, ReLU | (32, 128, 512) | (32, 128, 512) | (3,3) | 9,312 |
| **Output** |  |  |  |  |  |  |
| pred_class - $\hat{Y}$ | conv_op_0 | Conv2d, Softmax | (32, 128, 512) | $(C + 1, 128, 512)$ | (1,1) | $594 [(C + 1) \times 33]$ |
| pred_size - $\hat{S}$ | conv_op_0 | Conv2d, ReLU | (32, 128, 512) | (2, 128, 512) | (1,1) | 66 |
| **Total** |  |  |  |  |  | 1,890,388 |

### D MODEL ARCHITECTURE

This section provides additional details on the BatDetect2 model architecture, expanding upon the description in the main text. The model employs a 3-layer U-Net-style architecture (Ronneberger et al., 2015), incorporating an encoder, a decoder, and skip connections between them. A self-attention layer (Vaswani, 2017), denoted as `self_attn`, is incorporated in the central bottleneck of the model, enabling it to leverage information across extended timescales. This self-attention layer utilises a feature dimension of 256 and does not employ positional encoding. The model incorporates two specialised building blocks: `CoordConvDown` and `CoordConvUp`. The `CoordConvDown` layer performs the following sequence of operations: appending frequency coordinate information, 2D convolution, 2×2 max-pooling for downsampling, batch normalisation (BN) (Ioffe and Szegedy, 2015), followed by a ReLU non-linearity (Nair and Hinton, 2010). The `CoordConvUp` layer performs a similar, but inverse, set of operations, effectively upsampling the input tensor. This involves 2D bilinear upsampling, appending frequency coordinates, 2D convolution, batch normalisation, followed by a ReLU activation. The complete architecture of the BatDetect2 model is detailed in Table S6.

Following the model output, a non-maximal suppression operation is applied, implemented as two-dimensional max-pooling with a 9 × 9 kernel. The model then reports the top 200 events for each one-second segment of input audio, ranked by detection probability. Although the model can process input sequences of arbitrary length, in practice, it is recommended to segment longer audio files into clips of less than two seconds for independent processing.

### E TRAINING LOSSES

In this section we describe the training loss used by BatDetect2. The loss function is composed of three main terms and is inspired by those used in the CenterNet method for object detection in images (Zhou et al., 2019). The combined losses encourage the model to correctly predict the location, in frequency and time, of each echolocation call, the duration and frequency range of the call, and the species that is responsible for making the call.

Let us denote **x** ∈ ℝ^*H* ×*W*^ as the input spectrogram, with height *H* and width *W*. Here, height refers to the number of frequency bins and width is the number of temporal bins in the spectrogram. Prior to the final post-processing step (i.e., non-maximal suppression), the model outputs two tensors, *Ŷ* ∈ [0, 1]^*H* ×*W* ×(*C* +1)^ and 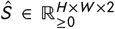. Here, *C* is the total number of species of interest, while the additional class is used to represent the background class (i.e., no bat present). *Ŷ* is the predicted species class probabilities and *Ŝ* contains the predicted size of any echolocation call estimated to be present. At training time the model has access to the ground truth values for *Y* and *S*. Both *Ŷ* and *Ŝ* contain an estimated value for each location in time and frequency space in the input spectrogram. For example, for a given frequency band *f* and time step *t, Ŝ*_*f t* 1_ encodes the predicted duration of the call (i.e., *t* _end_ − *t* _start_), and *Ŝ*_*f t* 2_ encodes the predicted frequency range of the call (i.e., *f*_max_ − *f*_min_). For a description of how *Y* and *S* are generated, please see the main text.

Additionally, let us define 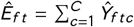, and similarly 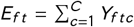 and *E* represent predicted and ground truth class-agnostic echolocation call scores, i.e., ‘Bat’ versus ‘Not bat’. Note that for *Ê* and *E*, the sum over does not include the background class. These additional terms are included as there are many instances in which the annotators have difficulty determining the correct species for a given call, and thus they can only label the event with the generic ‘Bat’ class label. This supervision can still be leveraged by allowing the model to determine which species may be present. The goal during training is to minimise the difference between the estimated *Ê, Ŷ*, and *Ŝ* and the respective ground truth values *E, Y*, and *S*. If successful, the model will be able to correctly predict the location in time and frequency of any echolocation call along with the species of the bat that generated the call.

#### E.1 Losses

The first loss encourages the model to correctly discriminate between bat echolocation calls and non-bat calls, i.e., background noise or other vocalising species. To achieve this, we use the focal loss (Lin et al., 2017), specifically, the keypoint variant of the focal loss from Law and Deng (2018), which is defined as:

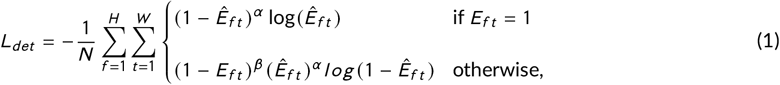

where *N* is the number of echolocation events in the spectrogram.

The next loss penalises the model for assigning the wrong species label to a detected echolocation call. This loss is similar *L*_*det*_, but instead of only discriminating between ‘Bat’ and ‘Not bat’, this loss encourages the model to predict the correct species label for each echolocation call:

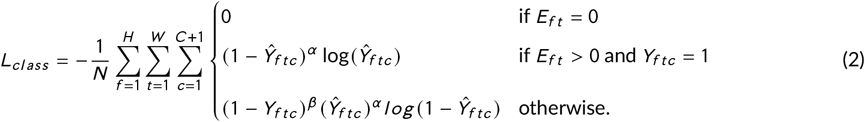

We use a masked version of the loss which is only applied to locations in the spectrogram where there is a echolocation call present, i.e., where *E*_*f t*_ > 0.

The final loss penalises the model for incorrectly predicting the ‘size’ of the predicted bounding box which overlaps with a ground truth echolocation call. Like *L*_*class*_, this loss is only applied to locations in time and frequency where an echolocation call in the training set has been annotated:

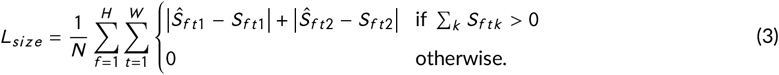

Here, ∑_*k*_ *S*_*f tk*_ > 0 simply indicates that this size loss is only applied where there is a echolocation call present. The final combined loss that is minimised during training is

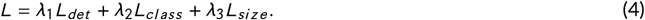

The loss is summed over each spectrogram in a given input training batch. During training, we set *λ*_1_, *λ*_2_, *λ*_3_ to 1.0, 2.0, and 0.1 respectively, and for both focal losses we set *α* = 2 and *β* = 4.

### F BASELINE MODEL

This appendix details the implementation and training procedures for the baseline models evaluated in this study.

#### F.1 Tadarida-D and Random Forest Baseline

For a given training dataset, we run Tadarida-D on each recording producing detected calls with extracted call features. These features are a set of 268 numerical values that encode information about the shape and frequency content of each detected call (see Bas et al., 2017 for further details).

To assign labels to each detected event, we compute the overlap between the event (using the reported time in file, duration, and frequency range from Tadarida-D) and the ground truth annotations. This overlap is quantified via the Intersection over Union (IoU) of the respective bounding boxes, with any overlaps below 0.01 being dropped. The predictions and ground truths are then matched using the `linear_sum_assignment` function from SciPy (Virtanen et al., 2020), which optimises the overall overlap across all potential pairings while ensuring each ground truth annotation is assigned to at most one predicted detection. Successfully matched events receive their corresponding species label, while any unmatched detection is assigned to the ‘Not bat’ class.

Finally we train a Random Forest (Breiman, 2001) classifier on the resulting features and labels. We use the implementation from `scikit-learn` (Pedregosa et al., 2011) with default parameters. We selected a Random Forest classifier to maintain consistency with the methodology of Bas et al. (2017), facilitating comparison of results.

#### F.2 Onset Detector Baseline

The Onset baseline serves as a simple, generic detector based on the acoustic strength envelope, implemented using the `onset_detect` function from the librosa library (McFee et al., 2015). To adapt the Onset detector for bat call detection, we performed hyperparameter optimisation using the Optuna framework (Akiba et al., 2019) to maximise the F1-score. The parameters optimised included `hop_length`, `delta`, `wait`, and `time_expansion`. Note that `time_expansion` is not part of the `onset_detect` function itself but was applied to the recordings prior to using the function. The optimisation was conducted over 40 trials using 200 random recordings sampled from the *UK*_same_ train set.

## References

Aide, T. M., C. Corrada-Bravo, M. Campos-Cerqueira, C. Milan, G. Vega, and R. Alvarez (2013). “Real-Time Bioacoustics Monitoring and Automated Species Identitication”. In: Peerj 1, e103. ISSN: 2167-8359. doi: 10.7717/peerj.103.

Akiba, T., S. Sano, T. Yanase, T. Ohta, and M. Koyama (2019). “Optuna: A Next-Generation Hyperparameter Optimization Framework”. In: The 25th ACM SIGKDD International Conference on Knowledge Dis-covery & Data Mining, pages 2623–2631.

Alipek, S., M. Maelzer, Y. Paumen, H. Schauer-Weisshahn, and J. Moll (2023). “An Efficient Neural Network Design Incorporating Autoencoders for the Classitication of Bat Echolocation Sounds”. In: Animals 13.16, page 2560. doi: 10.3390/ani13162560. PMID: 37627350.

Arias-Aguilar, A., F. Hintze, L. M. Aguiar, V. Rufray, E. Bernard, and M. J. R. Pereira (2018). “Who’s calling? Acoustic identitication of Brazilian bats”. In: Mammal Research 63.3, pages 231–253.

Bas, Y., D. Bas, and J.-F. Julien (2017). “Tadarida: A Tool-box for Animal Detection on Acoustic Recordings”. In: Journal of Open Research Software 5.1 (1), page 6. ISSN: 2049-9647. doi: 10.5334/jors.154.

Breiman, L. (2001). “Random Forests”. In: Machine Learning 45.1, pages 5–32. ISSN: 1573-0565. doi: 10.1023/A:1010933404324.

Brinkløv, S. M. M., J. Macaulay, C. Bergler, J. Tougaard, K. Beedholm, M. Elmeros, and P. T. Madsen (2023). “Open-Source Workflow Approaches to Passive Acoustic Monitoring of Bats”. In: Methods in Ecology and Evolution 14.7, pages 1747–1763. ISSN: 2041-210X. doi: 10.1111/2041-210X.14131.

Chen, X., J. Zhao, Y.-h. Chen, W. Zhou, and A. C. Hughes (2020). “Automatic Standardized Processing and Identitication of Tropical Bat Calls Using Deep Learning Approaches”. In: Biological Conservation 241, page 108269. ISSN: 0006-3207. doi: 10.1016/j.biocon.2019.108269.

Christin, S., É. Hervet, and N. Lecomte (2019). “Applications for Deep Learning in Ecology”. In: Methods in Ecology and Evolution 10.10, pages 1632–1644. doi: 10.1111/2041-210x.13256.

Denzinger, A. and H.-U. Schnitzler (2013). “Bat Guilds, a Concept to Classify the Highly Diverse Foraging and Echolocation Behaviors of Microchiropteran Bats”. In: Frontiers in Physiology 4. ISSN: 1664-042X. doi: 10.3389/fphys.2013.00164.

Dierckx, L., M. Beauvois, and S. Nijssen (2022). “Detection and Multi-label Classitication of Bats”. In: Advances in Intelligent Data Analysis XX. Edited by T. Bouadi, E. Fromont, and E. Hüllermeier. Lecture Notes in Computer Science. Cham: Springer International Publishing, pages 53–65. ISBN: 978-3-031-01333-1. doi: 10.1007/978-3-031-01333-1_5.

Dosovitskiy, A., L. Beyer, A. Kolesnikov, D. Weissenborn, X. Zhai, T. Unterthiner, M. Dehghani, M. Minderer, G. Heigold, S. Gelly, J. Uszkoreit, and N. Houlsby (2021). “An Image Is Worth 16×16 Words: Transformers for Image Recognition at Scale”. In: 2010.11929 [cs.CV]. doi: 10.48550/arXiv.2010.11929. arXiv: 2010.11929 [cs.CV].

Dumoulin, V., O. Stretcu, J. Hamer, L. Harrell, R. Laber, H. Larochelle, B. van Merriënboer, A. Navine, P. Hart, B. Williams, et al. (2025). “The search for squawk: Agile modeling in bioacoustics”. In: arXiv preprint 2505.03071.

Eichinski, P., C. Alexander, P. Roe, S. Parsons, and S. Fuller (2022). “A convolutional neural network bird species recognizer built from little data by iteratively training, detecting, and labeling”. In: Frontiers in Ecology and Evolution 10, page 810330.

Everingham, M., L. Van Gool, C. K. I. Williams, J. Winn, and A. Zisserman (2009). “The Pascal Visual Object Classes (VOC) Challenge”. In: International Journal of Computer Vision 88.2, pages 303–338. doi: 10.1007/s11263-009-0275-4.

Ferreira, D. F., R. Gibb, A. López-Baucells, N. J. Nunes, K. E. Jones, and R. Rocha (2022). “Species-Specitic Responses to Land-Use Change in Island Insectivorous Bats”. In: Journal for Nature Conservation 67, page 126177. doi: 10.1016/j.jnc.2022.126177.

Frick, W. F., T. Kingston, and J. Flanders (2019). “A Review of the Major Threats and Challenges to Global Bat Conservation”. In: Annals of the New York Academy of Sciences 1469.1, pages 5–25. ISSN: 1749-6632. doi: 10.1111/nyas.14045.

Fundel, F., D. A. Braun, and S. Gottwald (2023). “Automatic Bat Call Classitication Using Transformer Networks”. In: Ecological Informatics 78, page 102288. doi: 10.1016/j.ecoinf.2023.102288.

Gadot, T., S. Istrate, H. Kim, D. Morris, S. Beery, T. Birch, and J. Ahumada (2024). “To Crop or Not to Crop: Comparing Whole-image and Cropped Classitication on a Large Dataset of Camera Trap Images”. In: IET Computer Vision n/a.n/a. ISSN: 1751-9640. doi: 10.1049/cvi2.12318.

Gallacher, S., D. Wilson, A. Fairbrass, D. Turmukhambetov, M. Firman, S. Kreitmayer, O. Mac Aodha, G. Brostow, and K. Jones (2021). “Shazam for Bats: Internet of Things for Continuous Real-time Biodiversity Monitoring”. In: IET Smart Cities 3.3, pages 171–183. doi: 10.1049/smc2.12016.

Gibb, R., E. Browning, P. Glover-Kapfer, and K. E. Jones (2018). “Emerging Opportunities and Challenges for Passive Acoustics in Ecological Assessment and Monitoring”. In: Methods in Ecology and Evolution 10.2 (2), pages 169–185. ISSN: 2041-210X. doi: 10.1111/2041-210X.13101.

Hagiwara, M. (2023). “AVES: Animal Vocalization Encoder Based on Self-Supervision”. In: ICASSP 2023-2023 IEEE International Conference on Acoustics, Speech and Signal Processing (ICASSP), pages 1–5. doi: 10.1109/icassp49357.2023.10095642.

Hagiwara, M., B. Hoffman, J.-Y. Liu, M. Cusimano, F. Effenberger, and K. Zacarian (2023). “BEANS: The Benchmark of Animal Sounds”. In: ICASSP 2023 - 2023 IEEE International Conference on Acoustics, Speech and Signal Processing (ICASSP). ICASSP 2023 - 2023 IEEE International Conference on Acoustics, Speech and Signal Processing (ICASSP), pages 1–5. doi: 10.1109/ICASSP49357.2023.10096686.

He, K., X. Zhang, S. Ren, and J. Sun (2016). “Deep Residual Learning for Image Recognition”. In: Proceedings of the IEEE Conference on Computer Vision and Pattern Recognition, pages 770–778.

Heggan, C., S. Budgett, T. Hospedales, and M. Yaghoobi (2024). On the Transferability of Large-Scale Self-Supervision to Few-Shot Audio Classitication. doi: 10.48550/arXiv.2402.01274. 2402.01274 [cs]. url: http://arxiv.org/abs/2402.01274 (visited on 12/04/2024). Pre-published.

Hill, A. P., P. Prince, J. L. Snaddon, C. P. Doncaster, and A. Rogers (2019). “AudioMoth: A Low-Cost Acoustic Device for Monitoring Biodiversity and the Environment”. In: HardwareX 6, e00073. ISSN: 2468-0672. doi: 10.1016/j.ohx.2019.e00073.

Hoggatt, M. L., C. A. Starbuck, and J. M. O’Keefe (2024). “Acoustic Monitoring Yields Informative Bat Population Density Estimates”. In: Ecology and Evolution 14.2, e11051. ISSN: 2045-7758. doi: 10.1002/ece3.11051. PMID: 38389998.

Ioffe, S. and C. Szegedy (2015). “Batch Normalization: Accelerating Deep Network Training by Reducing Internal Covariate Shift”. 1502.03167 [cs].

Jones, G., D. S. Jacobs, T. H. Kunz, M. R. Willig, and P. A. Racey (2009). “Carpe Noctem: The Importance of Bats as Bioindicators”. In: Endangered Species Research 8.1–2, pages 93–115. ISSN: 1863-5407, 1613-4796. doi: 10.3354/esr00182.

Jones, G. and B. M. Siemers (2011). “The Communicative Potential of Bat Echolocation Pulses”. In: Journal of Comparative Physiology A 197.5, pages 447–457. ISSN: 1432-1351. doi: 10.1007/s00359-010-0565-x.

Jones, K. E., J. A. Russ, A.-T. Bashta, Z. Bilhari, C. Catto, I. Csősz, A. Gorbachev, P. Győrti, A. Hughes, I. Ivashkiv, N. Koryagina, A. Kurali, S. Langton, A. Collen, G. Margiean, I. Pandourski, S. Parsons, I. Prokofev, A. Szodoray-Paradi, F. Szodoray-Paradi, E. Tilova, C. L. Walters, A. Weatherill, and O. Zavarzin (2013). “Indicator Bats Program: A System for the Global Acoustic Monitoring of Bats”. In: Biodiversity Monitoring and Conservation. John Wiley & Sons, Ltd, pages 211–247. ISBN: 978-1-118-49074-7. doi: 10.1002/9781118490747.ch10.

Khalighifar, A., B. S. Gotthold, E. Adams, J. Barnett, L. O. Beard, E. R. Britzke, P. A. Burger, K. Chase, Z. Cordes, P. M. Cryan, E. Ferrall, C. T. Fill, S. E. Gibson, G. S. Haulton, K. M. Irvine, L. S. Katz, W. L. Kendall, C. A. Long, O. Mac Aodha, T. McBurney, S. McCarthy, M. W. McKown, J. O’Keefe, L. D. Patterson, K. A. Pitcher, M. Rustand, J. L. Segers, K. Seppanen, J. L. Siemers, C. Stratton, B. R. Straw, T. J. Weller, and B. E. Reichert (2022). “NABat ML: Utilizing Deep Learning to Enable Crowdsourced Development of Automated, Scalable Solutions for Documenting North American Bat Populations”. In: Journal of Applied Ecology 59.11, pages 2849–2862. ISSN: 1365-2664. doi: 10.1111/1365-2664.14280.

Kingma, D. P. and J. Ba (2017). “Adam: A Method for Stochastic Optimization”. 1412.6980 [cs].

Kobayashi, K., K. Masuda, C. Haga, T. Matsui, D. Fukui, and T. Machimura (2021). “Development of a Species Identitication System of Japanese Bats from Echolocation Calls Using Convolutional Neural Networks”. In: Ecological Informatics 62, page 101253. ISSN: 1574-9541. doi: 10.1016/j.ecoinf.2021.101253.

Lapp, S., T. Rhinehart, L. Freeland-Haynes, J. Khilnani, A. Syunkova, and J. Kitzes (2023). “OpenSoundscape: An Open-Source Bioacoustics Analysis Package for Python”. In: Methods in Ecology and Evolution 14.9, pages 2321–2328. ISSN: 2041-210X. doi: 10.1111/2041-210X.14196.

Law, H. and J. Deng (2018). “Cornernet: Detecting Objects as Paired Keypoints”. In: Proceedings of the European Conference on Computer Vision (ECCV), pages 734–750.

LeCun, Y., Y. Bengio, and G. Hinton (2015). “Deep Learning”. In: Nature 521.7553, pages 436–444. ISSN: 1476-4687. doi: 10.1038/nature14539.

Lin, T.-Y., P. Goyal, R. Girshick, K. He, and P. Dollar (2017). “Focal Loss for Dense Object Detection”. In: Proceedings of the IEEE International Conference on Computer Vision, pages 2980–2988.

Liu, R., J. Lehman, P. Molino, F. Petroski Such, E. Frank, A. Sergeev, and J. Yosinski (2018). “An Intriguing Failing of Convolutional Neural Networks and the Coord-Conv Solution”. In: Advances in Neural Information Processing Systems. Volume 31. Curran Associates, Inc.

Lostanlen, V., J. Salamon, M. Cartwright, B. McFee, A. Farnsworth, S. Kelling, and J. P. Bello (2019). “Per-Channel Energy Normalization: Why and How”. In: IEEE Signal Processing Letters 26.1, pages 39–43. ISSN: 1558-2361. doi: 10.1109/LSP.2018.2878620.

Mac Aodha, O., R. Gibb, K. E. Barlow, E. Browning, M. Firman, R. Freeman, B. Harder, L. Kinsey, G. R. Mead, S. E. Newson, I. Pandourski, S. Parsons, J. Russ, A. Szodoray-Paradi, F. Szodoray-Paradi, E. Tilova, M. Girolami, G. Brostow, and K. E. Jones (2018). “Bat Detective–Deep Learning Tools for Bat Acoustic Signal Detection”. In: PLOS Computational Biology 14.3, e1005995. ISSN: 1553-7358. doi: 10.1371/journal.pcbi.1005995.

MacSwiney G., M. C., F. M. Clarke, and P. A. Racey (2008). “What You See Is Not What You Get: The Role of Ultrasonic Detectors in Increasing Inventory Completeness in Neotropical Bat Assemblages”. In: Journal of Applied Ecology 45.5, pages 1364–1371. ISSN: 1365-2664. doi: 10.1111/j.1365-2664.2008.01531.x.

Mahbub, T., A. Bhagwagar, P. Chand, I. Zualkernan, J. Judas, and D. Dghaym (2024). “Bat2Web: a framework for real-time classitication of bat species echolocation signals using audio sensor data”. In: Sensors 24.9, page 2899.

Martínez Balvanera, S., O. Mac Aodha, M. J. Weldy, H. Pringle, E. Browning, and K. E. Jones (2024). “Whombat: An Open-source Audio Annotation Tool for Machine Learning Assisted Bioacoustics”. In: Methods in Ecology and Evolution n/a.n/a. ISSN: 2041-210X. doi: 10.1111/2041-210X.14468.

Martinez-de-Pison, F. J., J. Divasón, and F. González (2024). “BatNoiseDL: Discrimination of Bat Signals Through an Ensemble of Deep Learning Models”. In: International Conference on Hybrid Artiticial Intelligence Systems. Springer, pages 66–77.

McFee, B., C. Raffel, D. Liang, D. P. Ellis, M. McVicar, E. Battenberg, and O. Nieto (2015). “Librosa: Audio and Music Signal Analysis in Python.” In: SciPy, pages 18–24.

Menghani, G. (2023). “Efficient Deep Learning: A Survey on Making Deep Learning Models Smaller, Faster, and Better”. In: ACM Computing Surveys 55.12, pages 1–37. ISSN: 0360-0300, 1557-7341. doi: 10.1145/3578938. 2106.08962 [cs].

Meramo, K., P. Somervuo, M. Rannisto, P. Lauha, A. López-Baucells, F. Gili, T. Roslin, T. M. Lilley, and O. Ovaskainen (2025). “BSG-BATS: An open-source data annotation portal and classitier for European bat vocalizations”. In: Methods in Ecology and Evolution.

Mesaros, A., T. Heittola, T. Virtanen, and M. D. Plumbley (2021). “Sound Event Detection: A Tutorial”. In: IEEE Signal Processing Magazine 38.5, pages 67–83. ISSN: 1053-5888, 1558-0792. doi: 10.1109/MSP.2021.3090678. 2107.05463.

Montauban, C., M. Mas, C. Tuneu-Corral, O. S. Wangensteen, I. Budinski, J. Martí-Carreras, C. Flaquer, X. Puig-Montserrat, and A. López-Baucells (2021). “Bat Echolocation Plasticity in Allopatry: A Call for Caution in Acoustic Identitication of Pipistrellus Sp.” In: Behavioral Ecology and Sociobiology 75.4, page 70. ISSN: 1432-0762. doi: 10.1007/s00265-021-03002-7.

Nair, V. and G. E. Hinton (2010). “Rectitied Linear Units Improve Restricted Boltzmann Machines”. In: Proceedings of the 27th International Conference on Machine Learning (ICML-10), pages 807–814.

Obrist, M. and R. Boesch (2018). “BatScope Manages Acoustic Recordings, Analyses Calls, and Classities Bat Species Automatically”. In: Canadian Journal of Zoology 96.9, pages 939–954. doi: 10.1139/cjz-2017-0103.

Park, D. S., W. Chan, Y. Zhang, C.-C. Chiu, B. Zoph, E. D. Cubuk, and Q. V. Le (2019). SpecAugment: A Simple Data Augmentation Method for Automatic Speech Recognition. doi: 10.48550/arXiv.1904.08779. 1904.08779. url: http://arxiv.org/abs/1904.08779 (visited on 11/14/2024). Pre-published.

Parsons, S. and G. Jones (2000). “Acoustic Identitication of Twelve Species of Echolocating Bat By Discriminant Function Analysis and Artiticial Neural Networks”. In: Journal of Experimental Biology 203.17, pages 2641–2656. ISSN: 0022-0949. doi: 10.1242/jeb.203.17.2641.

Paszke, A., S. Gross, F. Massa, A. Lerer, J. Bradbury, G. Chanan, T. Killeen, Z. Lin, N. Gimelshein, L. Antiga Desmaison, A. Kopf, E. Yang, Z. DeVito, M. Raison, A. Tejani, S. Chilamkurthy, B. Steiner, L. Fang, J. Bai, and S. Chintala (2019). “PyTorch: An Imperative Style, High-Performance Deep Learning Library”. In: Advances in Neural Information Processing Systems 32. Curran Associates, Inc., pages 8024–8035.

Paumen, Y., M. Mälzer, S. Alipek, J. Moll, B. Lüdtke, and H. Schauer-Weisshahn (2021). “Development and Test of a Bat Calls Detection and Classitication Method Based on Convolutional Neural Networks”. In: Bioacoustics 31.5, pages 505–516. ISSN: 0952-4622. doi: 10.1080/09524622.2021.1978863.

Pedregosa, F., G. Varoquaux, A. Gramfort, V. Michel, Thirion, O. Grisel, M. Blondel, P. Prettenhofer, R. Weiss, V. Dubourg, J. Vanderplas, A. Passos, D. Cournapeau, M. Brucher, M. Perrot, and É. Duchesnay (2011). “Scikit-Learn: Machine Learning in Python”. In: Journal of Machine Learning Research 12.85, pages 2825–2830. ISSN: 1533-7928.

Pichler, M. and F. Hartig (2023). “Machine Learning and Deep Learning–A Review for Ecologists”. In: Methods in Ecology and Evolution 14.4, pages 994–1016. ISSN: 2041-210X. doi: 10.1111/2041-210X.14061.

Prat, Y., M. Taub, and Y. Yovel (2016). “Everyday Bat Vocalizations Contain Information about Emitter, Addressee, Context, and Behavior”. In: Scientitic Reports 6.1, page 39419. ISSN: 2045-2322. doi: 10.1038/srep39419.

Ravanelli, M. and Y. Bengio (2018). “Speaker Recognition from Raw Waveform with SincNet”. In: 2018 IEEE Spoken Language Technology Workshop (SLT). 2018 IEEE Spoken Language Technology Workshop (SLT), pages 1021–1028. doi: 10.1109/SLT.2018.8639585.

Redgwell, R. D., J. M. Szewczak, G. Jones, and S. Parsons (2009). “Classitication of echolocation calls from 14 species of bat by support vector machines and ensembles of neural networks”. In: Algorithms 2.3, pages 907–924.

Roemer, C., J.-F. Julien, P. P. Ahoudji, J.-M. Chassot, M. Genta, R. Colombo, G. Botto, C. A. Negreira, B. A. Djossa, R. K. Ing, A. Hassanin, V. Rufray, Q. Uriot, V.-C. Participants, and Y. Bas (2021). “An Automatic Classitier of Bat Sonotypes around the World”. In: Methods in Ecology and Evolution 12.12, pages 2432–2444. ISSN: 2041-210X. doi: 10.1111/2041-210X.13721.

Ronneberger, O., P. Fischer, and T. Brox (2015). “U-Net: Convolutional Networks for Biomedical Image Segmentation”. In: Medical Image Computing and Computer-Assisted Intervention – MICCAI 2015. Edited by N. Navab, J. Hornegger, W. M. Wells, and A. F. Frangi. Volume 9351. Cham: Springer International Publishing, pages 234–241. ISBN: 978-3-319-24573-7 978-3-319-24574-4. doi: 10.1007/978-3-319-24574-4_28.

Russo, D., L. Ancillotto, and G. Jones (2018). “Bats Are Still Not Birds in the Digital Era: Echolocation Call Variation and Why It Matters for Bat Species Identitication”. In: Canadian Journal of Zoology 96.2, pages 63–78. doi: 10.1139/cjz-2017-0089.

Russo, D., V. B. Salinas-Ramos, L. Cistrone, S. Smeraldo, L. Bosso, and L. Ancillotto (2021). “Do We Need to Use Bats as Bioindicators?” In: Biology 10.8, page 693. ISSN: 2079-7737. doi: 10.3390/biology10080693. PMID: 34439926.

Schwab, E., S. Pogrebnoj, M. Freund, F. Flossmann, S. Vogl, and K.-H. Frommolt (2022). “Automated Bat Call Classitication Using Deep Convolutional Neural Networks”. In: Bioacoustics 32.1, pages 1–16. ISSN: 0952-4622. doi: 10.1080/09524622.2022.2050816.

Sechidis, K., G. Tsoumakas, and I. Vlahavas (2011). “On the Stratitication of Multi-label Data”. In: Machine Learning and Knowledge Discovery in Databases. Edited by D. Gunopulos, T. Hofmann, D. Malerba, and M. Vazirgiannis. Volume 6913. Berlin, Heidelberg: Springer Berlin Heidelberg, pages 145–158. ISBN: 978-3-642-23807-9 978-3-642-23808-6. doi: 10.1007/978-3-642-23808-6_10.

Silva, B. and J. M. Herrera (2026). “A tine-tuning work-flow for CNN-based bat acoustic classitication”. In: Ecological Informatics, page 103645.

Simonyan, K. and A. Zisserman (2015). “Very Deep Convolutional Networks for Large-Scale Image Recognition”. 1409.1556 [cs].

Stowell, D. (2022). “Computational Bioacoustics with Deep Learning: A Review and Roadmap”. In: PeerJ 10, e13152. ISSN: 2167-8359. doi: 10.7717/peerj.13152.

Tabak, M. A., K. L. Murray, A. M. Reed, J. A. Lombardi, and K. J. Bay (2022). “Automated Classitication of Bat Echolocation Call Recordings with Artiticial Intelligence”. In: Ecological Informatics 68, page 101526. ISSN: 1574-9541. doi: 10.1016/j.ecoinf.2021.101526.

Van Merriënboer, B., J. Hamer, V. Dumoulin, E. Triantatillou, and T. Denton (2024). “Birds, Bats and beyond: Evaluating Generalization in Bioacoustics Models”. In: Frontiers in Bird Science 3. ISSN: 2813-3870. doi: 10.3389/fbirs.2024.1369756.

Vaswani, A. (2017). “Attention Is All You Need”. In: Advances in Neural Information Processing Systems.

Virtanen, P., R. Gommers, T. E. Oliphant, M. Haberland, T. Reddy, D. Cournapeau, E. Burovski, P. Peterson, W. Weckesser, J. Bright, S. J. van der Walt, M. Brett, J. Wilson, K. J. Millman, N. Mayorov, A. R. J. Nelson, E. Jones, R. Kern, E. Larson, C. J. Carey, İ. Polat, Y. Feng, E. W. Moore, J. VanderPlas, D. Laxalde, J. Perktold, R. Cimrman, I. Henriksen, E. A. Quintero, C. R. Harris, A. M. Archibald, A. H. Ribeiro, F. Pedregosa, and P. van Mulbregt (2020). “SciPy 1.0: Fundamental Algorithms for Scientitic Computing in Python”. In: Nature Methods 17.3 (3), pages 261–272. ISSN: 1548-7105. doi: 10.1038/s41592-019-0686-2.

Vogelbacher, M., H. Bellafkir, J. Gottwald, D. Schneider, M. Muhling, and B. Freisleben (2023). “Deep Learning for Recognizing Bat Species and Bat Behavior in Audio Recordings”. In: 2023 10th IEEE Swiss Conference on Data Science (SDS). 2023 10th IEEE Swiss Conference on Data Science (SDS), pages 50–57. doi: 10.1109/SDS57534.2023.00014.

Walters, C. L., R. Freeman, A. Collen, C. Dietz, M. Brock Fenton, G. Jones, M. K. Obrist, S. J. Puechmaille, T. Sattler, B. M. Siemers, et al. (2012). “A continentalscale tool for acoustic identitication of E uropean bats”. In: Journal of Applied Ecology 49.5, pages 1064–1074.

Walters, C. L., A. Collen, T. Lucas, K. Mroz, C. A. Sayer, and K. E. Jones (2013). “Challenges of Using Bioa-coustics to Globally Monitor Bats”. In: Bat Evolution, Ecology, and Conservation. Edited by R. A. Adams and S. C. Pedersen. New York, NY: Springer, pages 479–499. ISBN: 978-1-4614-7397-8. doi: 10.1007/978-1-4614-7397-8_23.

Wang, Y., P. Getreuer, T. Hughes, R. F. Lyon, and R. A. Saurous (2017). “Trainable Frontend for Robust and Far-Field Keyword Spotting”. In: 2017 IEEE International Conference on Acoustics, Speech and Signal Processing (ICASSP). 2017 IEEE International Conference on Acoustics, Speech and Signal Processing (ICASSP), pages 5670–5674. doi: 10.1109/ICASSP.2017.7953242.

YIlmaz, B., M. Sen, E. Masazade, and V. Beskardes (2022). “Behavior Classitication of Egyptian Fruit Bat (Rousettus Aegyptiacus) From Calls With Deep Learning”. In: Advances in Computational Intelligence and Robotics, pages 60–98. doi: 10.4018/978-1-7998-8686-0.ch004.

Zamora-Gutierrez, V., M. C. MacSwiney G. S. Martínez Balvanera, and E. Robredo Esquivelzeta (2021). “The Evolution of Acoustic Methods for the Study of Bats”. In: 50 Years of Bat Research: Foundations and New Frontiers. Edited by B. K. Lim, M. B. Fenton, R. M. Brigham, S. Mistry, A. Kurta, E. H. Gillam, A. Russell, and J. Ortega. Fascinating Life Sciences. Cham: Springer International Publishing, pages 43–59. ISBN: 978-3-030-54727-1. doi: 10.1007/978-3-030-54727-1_3.

Zhang, H., M. Cisse, Y. N. Dauphin, and D. Lopez-Paz (2018). “Mixup: Beyond Empirical Risk Minimization”. 1710.09412 [cs, stat].

Zhang, R. (2019). Making Convolutional Networks Shift-Invariant Again. doi: 10.48550/arXiv.1904.11486. 1904.11486 [cs]. url: http://arxiv.org/abs/1904.11486 (visited on 01/07/2025). Pre-published.

Zhou, X., D. Wang, and P. Krähenbühl (2019). “Objects as Points”. 1904.07850 [cs].

Zinck, R. (2023). BattyBirdNET - Bat Sound Analyzer. https://github.com/rdz-oss/BattyBirdNET-Analyzer.

Zualkernan, I., J. Judas, T. Mahbub, A. Bhagwagar, and P. Chand (2020). “A Tiny CNN Architecture for Identifying Bat Species from Echolocation Calls”. In: 2020 IEEE / ITU International Conference on Artiticial Intelligence for Good (AI4G). 2020 IEEE / ITU International Conference on Artiticial Intelligence for Good (AI4G), pages 81–86. doi: 10.1109/AI4G50087.2020.9311084.

